# Peripheral blood microbial signatures in COPD

**DOI:** 10.1101/2020.05.31.126367

**Authors:** Jarrett D. Morrow, Peter J. Castaldi, Robert P. Chase, Jeong H. Yun, Sool Lee, Yang-Yu Liu, Craig P. Hersh, the COPDGene Investigators

**Affiliations:** Channing Division of Network Medicine, Brigham and Women’s Hospital, Boston, MA; Division of Pulmonary and Critical Care Medicine, Brigham and Women’s Hospital, Boston, MA

**Keywords:** microbiome, COPD, emphysema, genomics, RNA-seq, transcriptomics

## Abstract

**Background:** The human microbiome has a role in the development of human diseases. Individual microbiome profiles are highly personalized, though many species are shared. Understanding the relationship between the human microbiome and disease may inform future individualized treatments. Specifically, the blood microbiome, once believed sterile, may be a surrogate for some lung and gut microbial characteristics. We sought associations between the blood microbiome and lung-relevant host factors.

**Methods:** Based on reads not mapped to the human genome, we detected microbial nucleic acid signatures in peripheral blood RNA-sequencing for 2,590 current and former smokers with and without chronic obstructive pulmonary disease (COPD) from the COPDGene study. We used the GATK microbial pipeline PathSeq to infer microbial profiles. We tested associations between the inferred profiles and lung disease relevant phenotypes and examined links to host gene expression pathways.

**Results:** The four phyla with highest abundance across all subjects were Proteobacteria, Actinobacteria, Firmicutes and Bacteroidetes. We observed associations between exacerbation phenotypes and the relative abundance of *Staphylococcus, Acidovorax* and *Cupriavidus*. The genus *Flavobacterium* was associated with emphysema and change in emphysema. Our host-microbiome interaction analysis revealed clustering of genera associated with emphysema, systemic inflammation, airway remodeling and exacerbations, through links to lung-relevant host pathways.

**Conclusions:** This study is the first to identify a bacterial microbiome signature in the peripheral blood of current and former smokers. Understanding the relationships between the systemic microbial populations and lung disease severity may inform novel interventions and aid in the understanding of exacerbation phenotypes.

## Introduction

The human microbiome has a role in human disease and overall health outcomes (1,2). Individual microbiome profiles are unique, although many species are shared (2,3). Knowledge of the relationship between the human microbial populations and disease may serve as a component of future comprehensive individualized treatment plans (4).

It was once believed that peripheral blood did not contain bacteria unless a serious infection was present. However, through use of culture-independent methods, evidence has emerged regarding the healthy human blood microbiome (5). The signature of the blood microbiome has provided insight into diseases such as schizophrenia (6), type 2 diabetes (7,8), chronic kidney disease (9) and liver fibrosis (10), and may provide a link to the microbiome in other tissues and diseases.

Relevance of the lung microbiome has been demonstrated in the context of lung diseases (12–14), including COPD (15,16), asthma (17,18) and IPF (19). In addition, exacerbations (20) and the healthy lung (21) have been explicitly addressed. These studies have involved both lung tissue (22,23) and the airway (24–26) with some researchers integrating the microbiome data with host gene expression (18,19,22,23,26). Study of the respiratory microbiome presents many challenges (27). Often at issue is the low biomass available in the samples (28). This is also an issue for peripheral blood microbiome studies (5).

Studies of the microbiome have typically involved 16S rRNA gene sequencing (29), with metagenomic sequencing emerging more recently (30). Repurposing read data from human sequencing studies that do not map to the human genome may reveal the microbiome, in parallel with the primary study. A possible application of this principle involves unmapped RNA-sequencing data (6,31,32), with detection via PathSeq (33), the microbial discovery pipeline for sequencing data available in the Genome Analysis Toolkit (GATK).

The foundation of this study is the detection of blood microbial signatures in RNA-sequencing data from a large subset of the COPDGene (Genetic Epidemiology of COPD) study. An overarching challenge in population based microbiome studies relates to statistical power, as testing for associations between the detected microbial profiles and variables of interest places demands on sample size. In our large analysis population, an enhanced power will enable findings in an environment such as blood, with its typically lower microbial signals. Using statistical tools for microbiome analysis, we tested associations between the detected taxa and a rich set of phenotype data in COPDGene. Network methods facilitated integration of the microbiome and host gene expression pathway data to highlight host-microbiome interactions. Previous use of the microbiome for diagnosis and prediction has proven successful in cancer (34), exacerbations of COPD (35), clinical outcomes in critically ill patients (36) and prediabetes (8). Our goal was to reveal microbial signatures in peripheral blood associated with lung disease relevant host factors and also to observe links with lung biology and microbiome through comparisons with prior studies. These signatures may serve as biomarkers of severity, decline, exacerbation susceptibility, or environmental exposure and may inform personalized treatment or diagnostic efforts. Some of the results of this study have been previously reported in the form of an abstract (37).

## Methods

### Study subjects

The analyses were performed on data from subjects enrolled in the COPDGene study (38) who participated in the five-year follow-up phase. COPDGene is a longitudinal cohort study that includes non-Hispanic Whites and African Americans enrolled at 21 centers across the United States. Study subjects provided written consent. COPDGene was approved by the Institutional Review Boards at all participating centers. The subjects include more than 10,000 current and former cigarette smokers with a minimum 10 pack-years smoking history, along with a small number of non-smokers. Cases have airflow obstruction (FEV1/FVC < 0.7) and control subjects had normal spirometry (FEV1% predicted ≥ 80% and FEV1/FVC ≥ 0.7). At the second phase follow-up visit, approximately five years after enrollment, a rich set of data were obtained through questionnaires, pre-and post-bronchodilator spirometry, volumetric computed tomography (CT) of the chest, and blood drawn for complete blood cell count, RNA-sequencing and biomarker studies. Emphysema severity was quantified via image analysis of chest CT data as the percentage of lung voxels below −950 HU (39).

### Microbial detection

We used reads not mapped to the human genome during the human gene expression analysis to detect a bacterial signature. Additional filtering of the unmapped reads was performed using the PathSeq microbial detection pipeline from the Genome Analysis Toolkit (GATK4) and the host reference available from the GATK Resource Bundle (33). This filtering addresses any remaining quality, host contamination or repetitive sequence issues. We subsequently used PathSeq to map these cleaned reads to bacterial genomes. The bacterial reference for this mapping was created using representative genomes, chromosomes, contigs and scaffolds (277,422 total genomic entries; September 25, 2019) from the National Center for Biotechnology Information (NCBI), and the PathSeq reference creation tools. Taxonomy information for these bacterial genomic data was also obtained from NCBI (RefSeq-release95.catalog.gz). Using these mapping results and taxonomy data, the inferred bacterial abundance profiles in each sample were assembled using PathSeq. Included in these profiling data were the raw read counts, adjusted scores and normalized scores (compositional data from the adjusted scores that represent inferred relative abundance) for taxa within each taxonomic classification (genera and phyla).

### Taxa associations

We tested associations between the normalized scores for each taxon and host variables using linear models with the R/Bioconductor package MaAsLin2 (Multivariate Association with Linear Models) (40). With relatively low levels of bacterial genetic content in peripheral blood, the data is inherently sparse and MaAsLin2 is particularly well suited for analysis of such microbial data. The base statistical model included the covariates age, sex, race, pack-years of smoking, current smoking status, and RNA-seq library preparation batch. For each of the models, adjustment for multiple testing controlled for false discovery rate (FDR < 5%). Scatter or box plots of the data for each model were produced by MaAsLin2.

### Contamination assessment

We sought to identify contamination (41) by testing the correlation between taxa abundance and RNA concentration, and observed possible contamination from the genus Methyloversatilis (rho < −0.4 and p < 0.0001; Figure E1 in the online supplement), a previously identified contaminant (42). Lacking a focus on diversity measures and detection of novel organisms in this study, the typical areas of impact for microbial contamination were minimized, and Methyloversatilis data were retained for later analysis. In addition, our analyses involve testing associations between host characteristics and microbial taxa, further muting the influence of batch-specific or study-wide contamination.

### Host microbe interactions

We projected the human gene expression data onto the pathways in the Hallmark gene set collection for the 3,304 genes at the intersection using gene set variation analysis via the R/Bioconductor package GSVA (43). The Hallmark canonical pathway set reduces redundancy found in public gene sets to enhance enrichment analyses. GSVA output is a pathway-by-subject matrix of expression data for observation of host-microbiome interactions. We used the pathways in this matrix as variables in MaAsLin2 models. We constructed a bipartite network (edges connecting taxa and pathways) using the results from these models. Communities within this network were identified using the R package CONDOR (44). Networks and communities were visualized using the R package igraph (45), with the Fruchterman-Reingold algorithm.

## Results

RNA-seq data were available for 2,647 samples from current and former smokers after exclusion of two samples with kinship issues. We performed microbial detection using PathSeq and excluded 57 samples with outlying unmapped read counts (see Methods). We then visualized the inferred relative abundance profiles and tested host associations (Figure 1) for these 2,590 subjects (Table 1). Ordered by mean normalized score from PathSeq, the four taxa observed at the phylum level above an abundance-filtering 1% threshold across all subjects were Proteobacteria, Actinobacteria, Firmicutes, and Bacteroidetes. In the abundance plot of the normalized scores for these four phyla, we observed batches with similar taxon distributions (Figure E2 in the online supplement). Twenty genera had mean normalized scores that eclipsed the 1% threshold. A heatmap was created for the normalized scores at the genus level, with clustering of samples in the columns by Bray-Curtis dissimilarity (Figure E3 in the online supplement). In the tracks for BMI, race, sex, batch, COPD status and smoking status, we observed visual clustering only by batch, justifying inclusion of this variable as a covariate in the statistical models.

**Figure 1.**
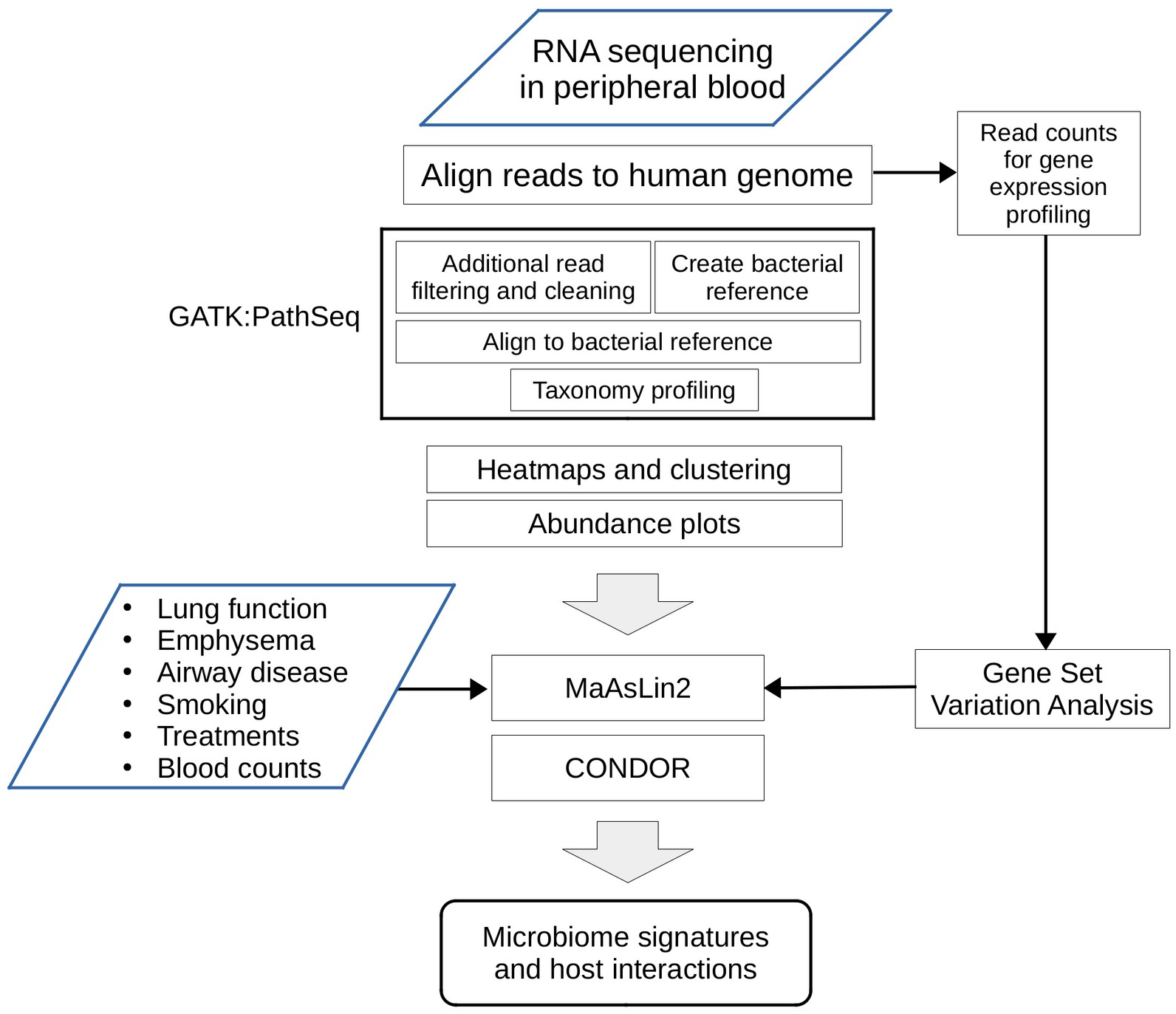
Overview of the study design illustrating the sequencing, statistical and gene enrichment framework. This illustrates the integration with host characteristics and gene expression for observations of host microbiome interaction (GATK = Genome Analysis Toolkit; MaAsLin2 = Multivariate Association with Linear Models)

**Table 1.** COPDGene study subjects

| **Demographics** | **N = 2590** |
| --- | --- |
|  | **Mean ± sd or distribution** |
| Age, years | 65.5 ± 8.6 |
| Sex (Female / Male) | 1257 / 1333 |
| Race (Non-Hispanic White / African American) | 1940 / 650 |
| Smoking (Current / Former) (n = 2580) | 909 / 1671 |
| Smoking History, pack-years (n = 2579) | 44.0 ± 24.0 |
| GOLD status (n = 2541) |  |
| 4 | 101 |
| 3 | 245 |
| 2 | 505 |
| 1 | 258 |
| 0 (controls) | 1101 |
| -1 (PRISm) | 331 |
| FEV 1 % predicted (n = 2541) | 78.6 ± 24.2 |
| FEV 1 / FVC (n = 2540) | 0.68 ± 0.15 |
| Percent emphysema at -950HU (n = 2388) | 5.5 ± 9.2 |
| Body mass index kg/m<sup>2</sup> (n = 2581) | 29.0 ± 6.3 |
| Airway wall thickness, segmental bronchi (n = 2385) | 1.03 ± 0.22 |
| Severe Exacerbation in the year prior (no / yes) (n = 2581) | 2367 / 214 |
| Treated with Oral Corticosteroids (no / yes) (n = 2538) | 2504 / 34 |
| Survival (alive / deceased) * | 2431 / 159 |
| Abbreviations: FEV1=forced expiratory volume in 1 sec; FVC= forced vital capacity; PRISm = Preserved Ratio Impaired Spirometry (FEV1<80% predicted with FEV1/FVC≥0.7) | |
| * survival status as of October 2018 |  |

### Genera abundance and host phenotype

We tested associations between the inferred relative abundance for each taxon at the genus level and host phenotype, exposure, treatment and trait variables using linear models with MaAsLin2 (Table E1 in the online supplement). We summarized the findings (Tables E2-E32 in the online supplement) in a heatmap of the q-values and effect sizes for each test (Figure 2). Scatter or box plots for the significant (FDR < 5%) findings illustrate the relationships between inferred taxa abundance and the variable of interest (Figures E4-E33 in the online supplement). We observed significant associations between the genus *Cupriavidus* and both exacerbation frequency (q = 0.022) and the history of a severe exacerbations in the prior year (q = 0.0077). *Staphylococcus* and *Acidovorax* genera were also associated with severe exacerbations (q = 0.025 and q = 0.021, respectively). Airway wall thickness on quantitative analysis of chest CT scans was associated with *Staphylococcus* abundance (q = 0.0028). Inferred abundance of the genus *Flavobacterium* was significantly associated with higher percent emphysema on chest CT scan (q = 0.0054) and an increase in percent emphysema from enrollment to second phase follow-up (q = 0.04). Abundance of *Flavobacterium* was also associated with reduced lymphocyte count (q = 0.026). For the case-control analysis, *Streptococcus* abundance was associated with COPD case-control status (q = 0.043). We also observed an association between *Streptococcus* and oral steroid use (q = 0.037). *Methylobacterium* abundance was significantly associated with years since quitting cigarette smoking (q = 0.015) and pack-years history of smoking (q = 0.028) and *Methylorubrum* abundance was associated with pack-years (q = 0.047). *Methyloversatilis* abundance was associated with current smoking status (q = 0.0092). A significant association was observed for the genus *Nevskia* and current smoking status (q = 0.023).

**Figure 2.**
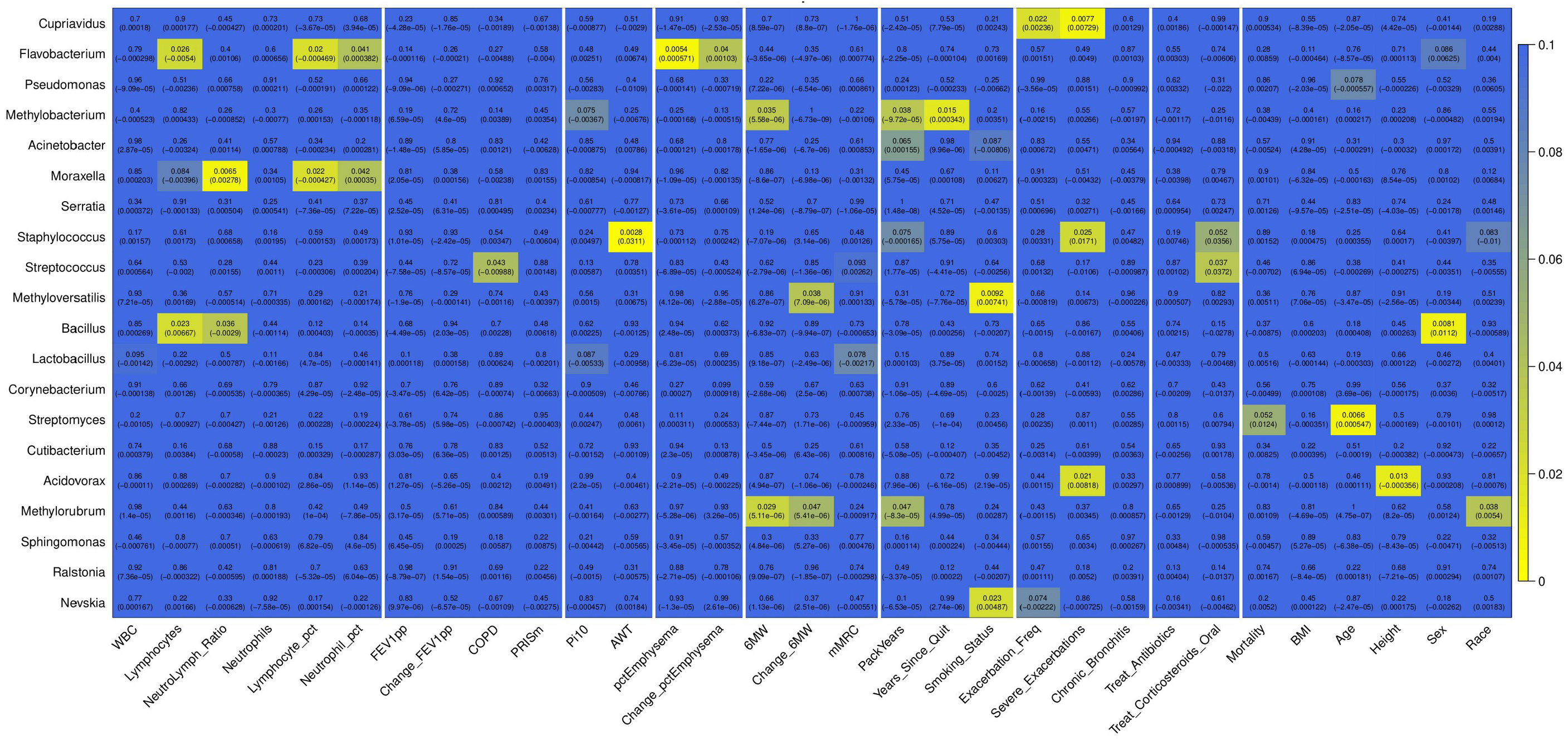
Heatmap of the associations between genera inferred relative abundance and host-related variables. Top entry in each cell is the q-value (FDR adjusted p-value for each variable) and the bottom is the effect estimate from the MaAsLin2 model. Significant findings have q-values less than 0.05.

### Phyla abundance and host phenotype

We also tested associations between the inferred relative abundance for each taxon at the phylum level using the same methods and summarized the findings in a heatmap (Figure E34 in the online supplement); we observed at least one significant (FDR < 5%) association for two of the four phyla: Bacteroidetes and Proteobacteria. We observed significant associations between Bacteroidetes inferred abundance and lymphocyte count (q = 0.018), change in FEV1 percent predicted from Phase 1 to Phase 2 (q = 0.0063) and percent emphysema (q = 0.013). We also observed associations between Proteobacteria abundance and neutrophil percentage (q = 0.026), severe exacerbations (q = 0.047) and race (q = 0.045).

### Host-microbiome interactions

We sought to highlight host-microbiome interactions using microbial abundance profiles and host gene expression pathway data. We created a matrix of pathway expression for the Hallmark sets from MSigDB using the R/Bioconductor package GSVA and the host gene expression data. We tested the association between taxa and host pathways for each of the 20 genera using models, adjusting for age, sex, race, pack-years of smoking, current smoking status, and library prep batch. The associations across all taxa and pathways were summarized in a heatmap (Figure E35 in the online supplement). We used network methods to visualize the large set of significant (FDR < 5%) findings. We constructed a bipartite network using the significant associations as edges between taxa and pathways (Figure E36 in the online supplement). Using CONDOR, we identified five communities within this network (Figures E37 – E40 in the online supplement) with one of particular relevance to lung disease (Figure 3). This community has five genera (*Bacillus*, *Flavobacterium*, *Acinetobacter*, *Staphylococcus* and *Acidovorax*) and 11 host pathways. Within these communities we observe clustering of genera with shared pathway associations, suggesting joint influence on the host processes.

**Figure 3.**
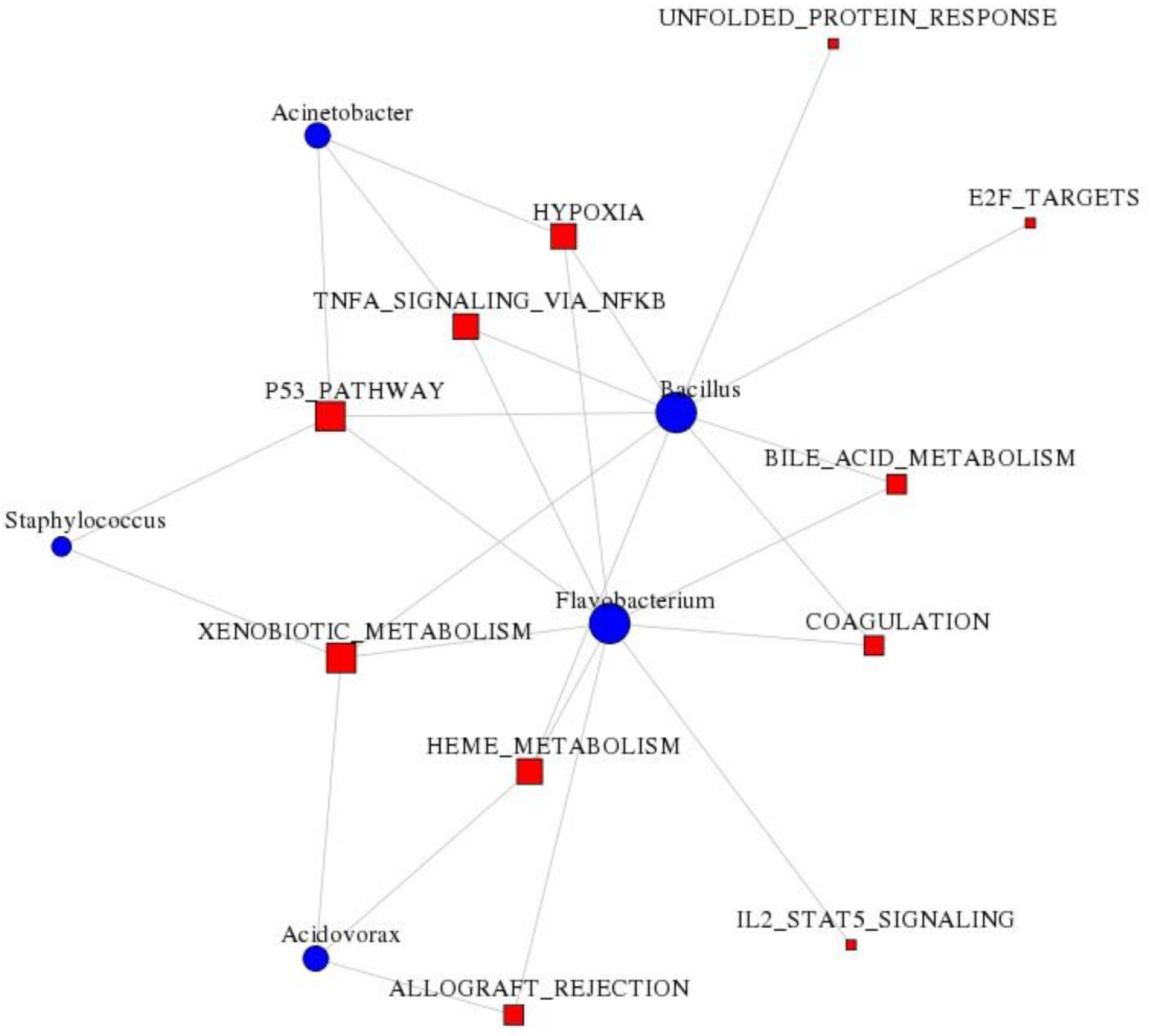
Community from the bipartite network from the host-microbiome interaction analysis with relevance to inflammation, emphysema, exacerbation severity and airway disease. Edges represent a significant (FDR < 5%) association between inferred genus abundance and the expression of the Hallmark pathway in the human host. The red squares represent Hallmark pathways from MSigDB and the blue circles represent genera.

## Discussion

We profiled the peripheral blood microbiome in COPDGene Study participants using the RNA-sequencing reads that did not map to the human genome and detected taxa at both the phylum and genus levels. We tested associations between inferred taxa abundance and host-related variables. At the phylum level, we identified Proteobacteria, Actinobacteria, Firmicutes, and Bacteroidetes. Once believed sterile, studies have shown healthy peripheral blood microbiomes typically include a nucleic acid signature of these phyla (5,32,46). Detection at the genus level produced a larger set of taxa, many of which (12 of 20) had inferred abundance significantly associated (FDR < 5%) with at least one host-related variable. Nine of the genera had more than one significant finding.

### Phylum associations

Both Bacteroidetes and Proteobacteria have been identified in lung tissue (14,26,47,48), with Bacteroidetes relative abundance lower in COPD (23,48). Proteobacteria relative abundance has been found to be higher in sputum during exacerbations (26,48,49) and negatively correlated with neutrophil infiltration (23). Our Proteobacteria findings for severe exacerbations are consistent with these previous studies in the lung. However, the neutrophil-percentage finding for Proteobacteria, and the Bacteroidetes results for percent emphysema and lung function decline, have opposite effect compared with findings in lung tissue. Further studies will be required to determine whether these associations highlight cross-tissue mechanisms supportive or similar to the immunomodulatory gut-lung axis (50), or perhaps ones analogous to liver disease and gut microbiome interactions and translocation events (51). Despite the direction of effect differences, together these findings suggest we may be capturing lung disease relevant microbial signatures at the phylum level in peripheral blood.

### Exacerbations and emphysema

We observed associations of the genus *Cupriavidus* with both exacerbation frequency and a recent history of severe exacerbations. Species within *Cupriavidus* have been recovered from respiratory tract specimens in Cystic Fibrosis patients (52,53), although the extent of their role in exacerbations is not clear. We found *Staphylococcus* and *Acidovorax* to be associated with exacerbations, and both species have been previously identified in the airway of COPD patients treated for severe exacerbations (54), with *Staphylococcus* species previously suggested to have a role in exacerbations of COPD (55,56) and chronic airway inflammation (57). Recurrent exacerbations are expected to have an impact on airway dimensions. However, in contrast to this observed association of *Staphylococcus* and airway wall thickness, the presence of bacteria in the lower airway in asthma did not previously have an observed impact on airway wall thickness (58). Species in the genus *Flavobacterium* have been found to be negatively correlated with CD4+ T cells and positively correlated with alveolar surface area in lung tissue (23). Our lymphocyte finding is concordant with these lung results. For the case-control analysis, *Streptococcus* abundance was associated with COPD status. *Streptococcus pneumoniae* is a common cause of respiratory infections and has been isolated from sputum samples from a population of COPD patients in both a stable and an exacerbation state (59).

### Smoking and inflammation

*Methylobacterium*, *Methylorubrum* and *Methyloversatilis* abundances were associated with smoking-related measures. The methylotrophic nature of species within these three genera (phylum Proteobacteria) suggests a role in degrading chemicals produced by cigarette smoke (60). *Methylobacterium* also has a putative role in xenobiotic metabolism in the gut microbiome (61). Last, abundance of Proteobacteria in human intestine has been found to change with smoking cessation (62). *Moraxella* and *Bacillus* abundance were significantly associated with lymphocytes or neutrophils, and although we did not observe significant associations with lung-related phenotypes for these in this study, they both have putative roles in COPD and airway inflammation (63,64). The association of both *Moraxella* and *Bacillus* with the ratio of neutrophils to lymphocytes highlights a systemic inflammatory response, analogous to observations in the gut microbiome (65).

### Pathways group taxa within host-microbiome networks

We explored host-microbiome interactions using network methods for significant taxa and host pathway associations. Within the communities of the bipartite network, genera with common pathway associations were clustered, providing insight into shared influence on the host processes. For one particular community (Figure 3) within the bipartite network, we observed clustering of *Flavobacterium* (associated with emphysema severity) with *Bacillus* (associated with systemic inflammation), through host xenobiotic metabolism, P53 tumor suppressor, Tumor Necrosis Factor (TNF)-alpha (TNFA), heme metabolism, hypoxia and coagulation pathways. This community also links the genera *Staphylococcus* and *Acidovorax* (airway remodeling and exacerbation associated) through a subset of these pathways. These pathways are involved in aspects of lung disease. Heme metabolism is relevant in COPD; abnormal iron regulation contributes to cellular perturbations in lung tissue (66). TNF-alpha has been implicated as a biomarker and possible therapeutic target for COPD (67). Alveolar hypoxia increases with disease severity in COPD and uncorrected chronic hypoxemia is associated with adverse outcomes (68). This bipartite network approach demonstrates a versatile method for observation of host-microbiome interactions, both within and across host tissues.

There are several limitations to the current study. Using RNA-sequencing data, we are capturing RNA from bacterial gene expression. However, we did not seek a gene-level analysis. Instead, these mapped reads are serving as a proxy for abundance. Variation in gene expression activity would therefore impact our estimations of taxa abundance (22), as would the presence of RNA from bacterial expression in other tissues or compartments. Future studies involving 16S rRNA gene or whole-genome shotgun sequencing in parallel with the host transcriptome analysis will address this limitation. In the case of survival status, we applied a linear model through MaAsLin2 instead of a Cox model, that would have incorporated time to event information. Observing concordance of the blood microbiome with other tissues will be challenging, though cross-tissue studies are possible (11), and previous studies have linked peripheral blood metabolome to the gut microbiome (69). Lack of concordance or direction of effect does not entirely diminish the role of these findings with respect to biomarkers, and future metagenomic studies will seek to analyze data across tissues from the same subjects.

In this study of the blood microbiome, we were able to detect lung disease relevant bacterial signatures in peripheral blood RNA-seq data from a large cohort of smokers. Analyses at the genus level provided insight into variation in the blood microbiome associated with host-related factors. Using a network approach, we identified host transcriptomic pathways linking multiple taxa, highlighting a useful method for future studies of the human microbiome and transcriptome. Together these findings demonstrate that the peripheral blood microbiome has the potential to capture relevant lung microbiome features and biology, and perhaps provide a foundation for discovery of blood markers for personalized treatments and predictive disease models.

## COPDGene Phase 3

### Grant Support and Disclaimer

The project described was supported by Award Number U01 HL089897 and Award Number U01 HL089856 from the National Heart, Lung, and Blood Institute. The content is solely the responsibility of the authors and does not necessarily represent the official views of the National Heart, Lung, and Blood Institute or the National Institutes of Health.

### COPD Foundation Funding

COPDGene is also supported by the COPD Foundation through contributions made to an Industry Advisory Board comprised of AstraZeneca, Boehringer-Ingelheim, Genentech, GlaxoSmithKline, Novartis, Pfizer, Siemens, and Sunovion.

### COPDGene^®^ Investigators – Core Units

*Administrative Center*: James D. Crapo, MD (PI); Edwin K. Silverman, MD, PhD (PI); Barry J. Make, MD; Elizabeth A. Regan, MD, PhD

*Genetic Analysis Center*: Terri Beaty, PhD; Ferdouse Begum, PhD; Peter J. Castaldi, MD, MSc; Michael Cho, MD; Dawn L. DeMeo, MD, MPH; Adel R. Boueiz, MD; Marilyn G. Foreman, MD, MS; Eitan Halper-Stromberg; Lystra P. Hayden, MD, MMSc; Craig P. Hersh, MD, MPH; Jacqueline Hetmanski, MS, MPH; Brian D. Hobbs, MD; John E. Hokanson, MPH, PhD; Nan Laird, PhD; Christoph Lange, PhD; Sharon M. Lutz, PhD; Merry-Lynn McDonald, PhD; Margaret M. Parker, PhD; Dmitry Prokopenko, Ph.D; Dandi Qiao, PhD; Elizabeth A. Regan, MD, PhD; Phuwanat Sakornsakolpat, MD; Edwin K. Silverman, MD, PhD; Emily S. Wan, MD; Sungho Won, PhD

*Imaging Center*: Juan Pablo Centeno; Jean-Paul Charbonnier, PhD; Harvey O. Coxson, PhD; Craig J. Galban, PhD; MeiLan K. Han, MD, MS; Eric A. Hoffman, Stephen Humphries, PhD; Francine L. Jacobson, MD, MPH; Philip F. Judy, PhD; Ella A. Kazerooni, MD; Alex Kluiber; David A. Lynch, MB; Pietro Nardelli, PhD; John D. Newell, Jr., MD; Aleena Notary; Andrea Oh, MD; Elizabeth A. Regan, MD, PhD; James C. Ross, PhD; Raul San Jose Estepar, PhD; Joyce Schroeder, MD; Jered Sieren; Berend C. Stoel, PhD; Juerg Tschirren, PhD; Edwin Van Beek, MD, PhD; Bram van Ginneken, PhD; Eva van Rikxoort, PhD; Gonzalo Vegas Sanchez-Ferrero, PhD; Lucas Veitel; George R. Washko, MD; Carla G. Wilson, MS;

*PFT QA Center, Salt Lake City, UT*: Robert Jensen, PhD

*Data Coordinating Center and Biostatistics, National Jewish Health, Denver, CO*: Douglas Everett, PhD; Jim Crooks, PhD; Katherine Pratte, PhD; Matt Strand, PhD; Carla G. Wilson, MS

*Epidemiology Core, University of Colorado Anschutz Medical Campus, Aurora, CO*: John E. Hokanson, MPH, PhD; Gregory Kinney, MPH, PhD; Sharon M. Lutz, PhD; Kendra A. Young, PhD

*Mortality Adjudication Core:* Surya P. Bhatt, MD; Jessica Bon, MD; Alejandro A. Diaz, MD, MPH; MeiLan K. Han, MD, MS; Barry Make, MD; Susan Murray, ScD; Elizabeth Regan, MD; Xavier Soler, MD; Carla G. Wilson, MS

*Biomarker Core*: Russell P. Bowler, MD, PhD; Katerina Kechris, PhD; Farnoush Banaei-Kashani, Ph.D

### COPDGene^®^ Investigators – Clinical Centers

*Ann Arbor VA:* Jeffrey L. Curtis, MD; Perry G. Pernicano, MD

*Baylor College of Medicine, Houston, TX*: Nicola Hanania, MD, MS; Mustafa Atik, MD; Aladin Boriek, PhD; Kalpatha Guntupalli, MD; Elizabeth Guy, MD; Amit Parulekar, MD;

*Brigham and Women’s Hospital, Boston, MA*: Dawn L. DeMeo, MD, MPH; Alejandro A. Diaz, MD, MPH; Lystra P. Hayden, MD; Brian D. Hobbs, MD; Craig Hersh, MD, MPH; Francine L. Jacobson, MD, MPH; George Washko, MD

*Columbia University, New York, NY*: R. Graham Barr, MD, DrPH; John Austin, MD; Belinda D’Souza, MD; Byron Thomashow, MD

*Duke University Medical Center, Durham, NC*: Neil MacIntyre, Jr., MD; H. Page McAdams, MD; Lacey Washington, MD

*Grady Memorial Hospital, Atlanta, GA*: Eric Flenaugh, MD; Silanth Terpenning, MD

*HealthPartners Research Institute, Minneapolis, MN*: Charlene McEvoy, MD, MPH; Joseph Tashjian, MD

*Johns Hopkins University, Baltimore, MD*: Robert Wise, MD; Robert Brown, MD; Nadia N. Hansel, MD, MPH; Karen Horton, MD; Allison Lambert, MD, MHS; Nirupama Putcha, MD, MHS

*Lundquist Institute for Biomedical Innovationat Harbor UCLA Medical Center, Torrance, CA*: Richard Casaburi, PhD, MD; Alessandra Adami, PhD; Matthew Budoff, MD; Hans Fischer, MD; Janos Porszasz, MD, PhD; Harry Rossiter, PhD; William Stringer, MD

*Michael E. DeBakey VAMC, Houston*, *TX*: Amir Sharafkhaneh, MD, PhD; Charlie Lan, DO

*Minneapolis VA:* Christine Wendt, MD; Brian Bell, MD; Ken M. Kunisaki, MD, MS

*National Jewish Health, Denver, CO*: Russell Bowler, MD, PhD; David A. Lynch, MB

*Reliant Medical Group, Worcester, MA*: Richard Rosiello, MD; David Pace, MD

*Temple University, Philadelphia, PA:* Gerard Criner, MD; David Ciccolella, MD; Francis Cordova, MD; Chandra Dass, MD; Gilbert D’Alonzo, DO; Parag Desai, MD; Michael Jacobs, PharmD; Steven Kelsen, MD, PhD; Victor Kim, MD; A. James Mamary, MD; Nathaniel Marchetti, DO; Aditi Satti, MD; Kartik Shenoy, MD; Robert M. Steiner, MD; Alex Swift, MD; Irene Swift, MD; Maria Elena Vega-Sanchez, MD

*University of Alabama, Birmingham, AL:* Mark Dransfield, MD; William Bailey, MD; Surya P. Bhatt, MD; Anand Iyer, MD; Hrudaya Nath, MD; J. Michael Wells, MD

*University of California, San Diego, CA*: Douglas Conrad, MD; Xavier Soler, MD, PhD; Andrew Yen, MD

*University of Iowa, Iowa City, IA*: Alejandro P. Comellas, MD; Karin F. Hoth, PhD; John Newell, Jr., MD; Brad Thompson, MD

*University of Michigan, Ann Arbor, MI:* MeiLan K. Han, MD MS; Ella Kazerooni, MD MS; Wassim Labaki, MD MS; Craig Galban, PhD; Dharshan Vummidi, MD

*University of Minnesota, Minneapolis, MN*: Joanne Billings, MD; Abbie Begnaud, MD; Tadashi Allen, MD

*University of Pittsburgh, Pittsburgh, PA*: Frank Sciurba, MD; Jessica Bon, MD; Divay Chandra, MD, MSc; Carl Fuhrman, MD; Joel Weissfeld, MD, MPH

*University of Texas Health, San Antonio, San Antonio, TX*: Antonio Anzueto, MD; Sandra Adams, MD; Diego Maselli-Caceres, MD; Mario E. Ruiz, MD; Harjinder Singh

