## Supplemental Material for "Peripheral blood microbial signatures in COPD"

### Supplemental Methods

RNA sequencing

Sample processing was previously described for earlier stages of these data (1) and those data are available in GEO (accession number GSE97531). Briefly, total RNA was extracted from PAXgene™ Blood RNA tubes using the Qiagen PreAnalytiX PAXgene Blood miRNA Kit (Qiagen, Valencia, CA). Extracted RNA samples with RNA integrity number greater than seven and a concentration of at least 25 µg/ul were sequenced. Globin reduction and cDNA library preparation for total RNA was performed with the Illumina TruSeq Stranded Total RNA with Ribo-Zero Globin kit (Illumina, Inc., San Diego, CA). Paired end reads with nominal 75 bp length were generated on an Illumina HiSeq 2500 flow cell. Sequencing was performed to an average depth of 20 million reads.

Data Processing

The quality control pipeline for these sequencing reads included FastQC (2) and RNA-SeQC (3). Adapter trimming was performed using Skewer (4). STAR aligner version 2.4.0 h (5) was used to map the reads to the GRCH38 genome reference and RSubreads produced gene-level counts (6) with the Ensembl version 81 gene annotation (7). As part of the cleaning and quality control process, we confirmed expression consistent with reported sex, and concordance between variants called from RNA sequencing reads and corresponding DNA genotyping. Two samples were excluded due to kinship issues. Data for genes with variance in the upper 90th percentile and average read counts greater than five were retained and intersected with the Hallmark gene sets from MSigDB (8). A total of 3,304 genes were included in the host interaction analysis.

Microbial detection – quality control

During quality evaluation, we removed one outlying processing batch (57 samples) with a mean total read count four standard deviations from the mean of the total read data. Heatmaps of taxa and samples were produced using the R package pheatmap (9) with visual clustering of samples performed using Bray-Curtis dissimilarity from the vegdist function in the R package vegan (10) and clustering of taxa by euclidean distance. Abundance plots were created using the R package ggplot2 (11).

**Supplemental Figures**

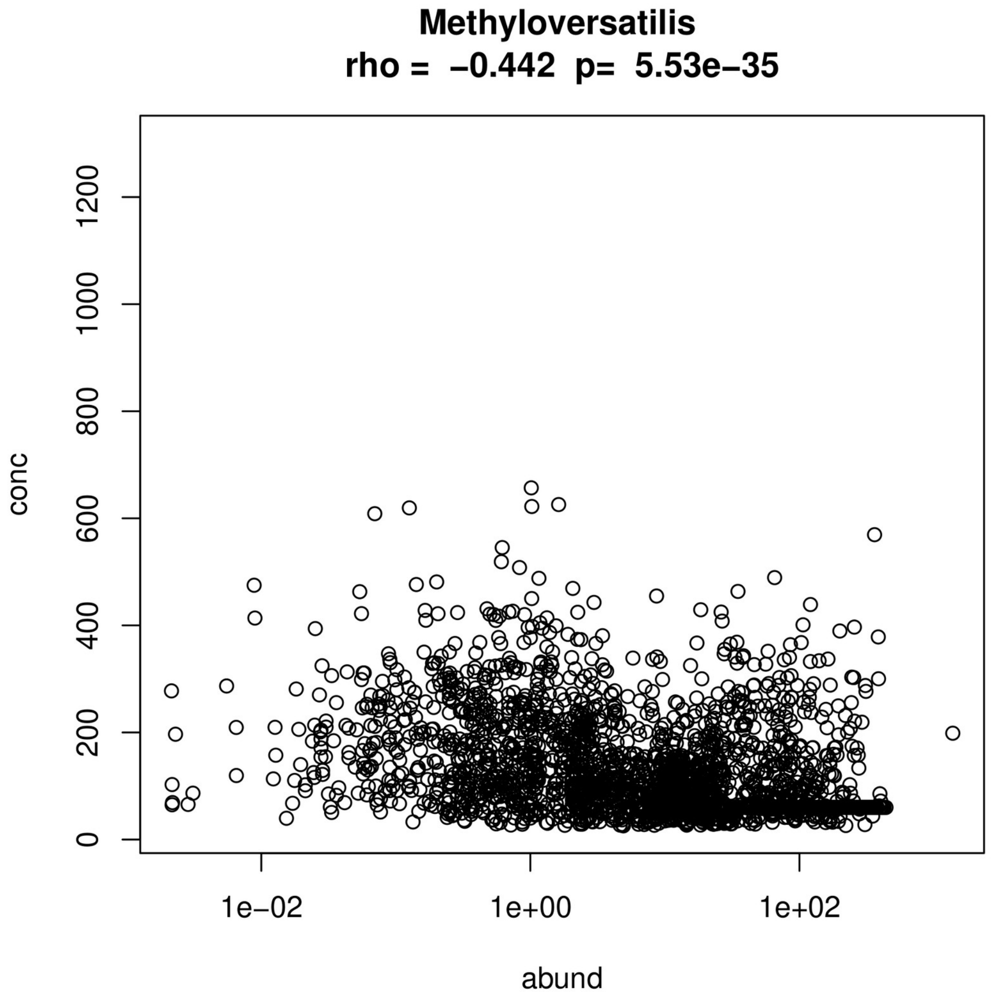

Figure E1. Scatter plot of abundance and DNA concentration for Methyloversatilis.

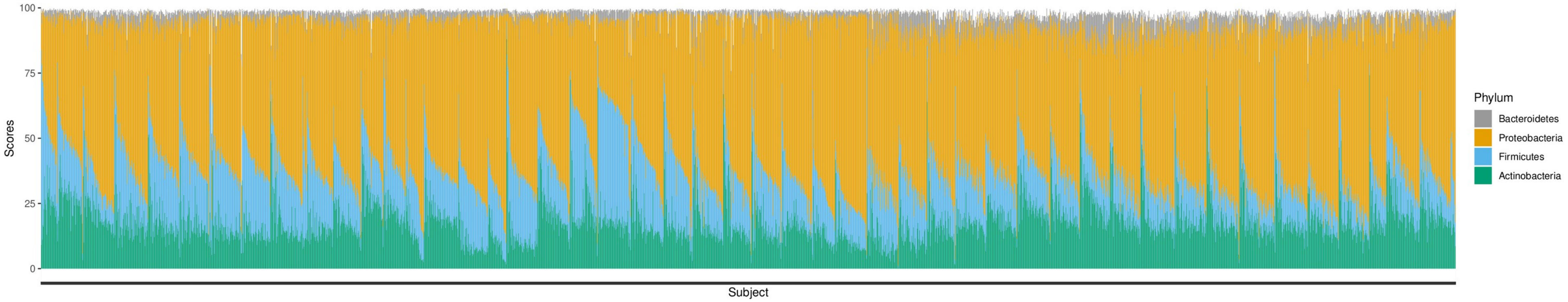

Figure E2. Abundance plots of the normalized scores for the top four phyla

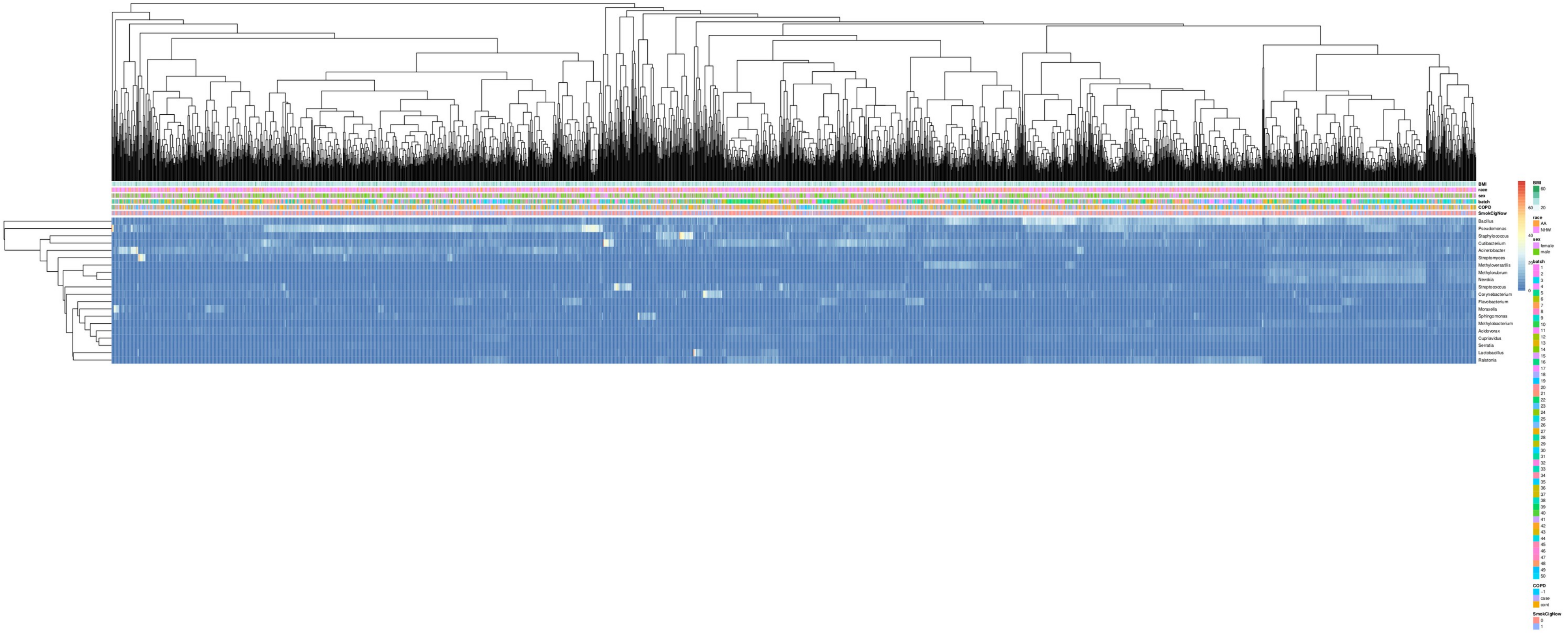

Figure E3. Heatmap of the normalized scores at the genus level, with clustering of samples in the columns by Bray-Curtis dissimilarity. Tracks are included for BMI, race, sex, batch, COPD status and smoking status.

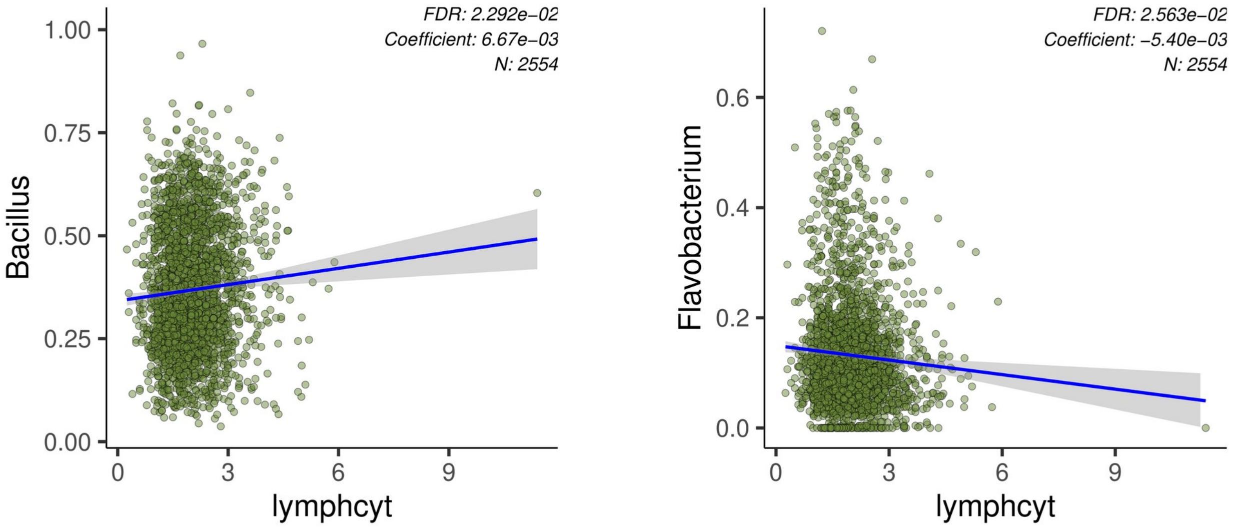

Figures E4 and E5. Plots of inferred taxa abundance and lymphocyte count

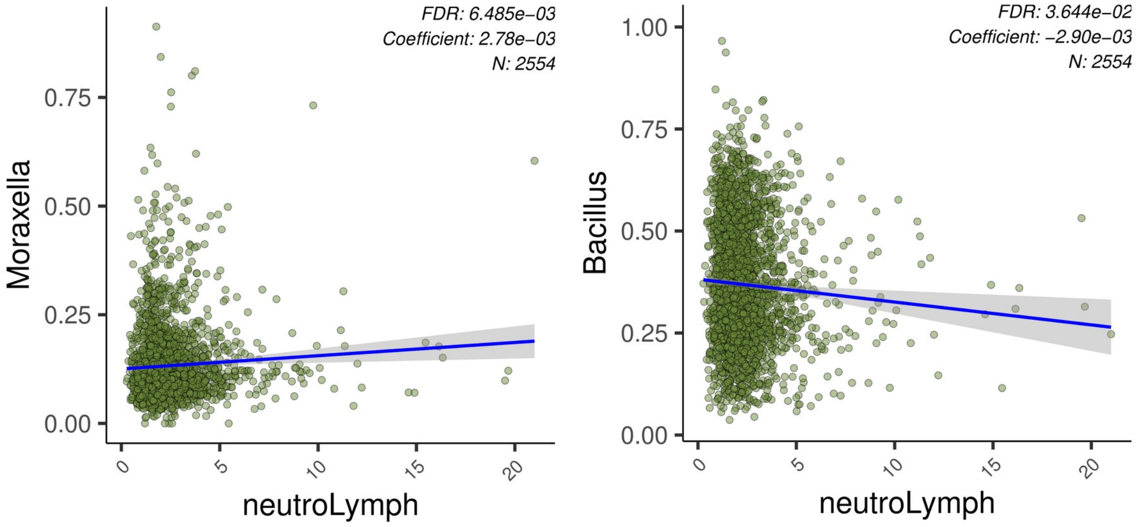

Figures E6 and E7. Plots of inferred taxa abundance and neutrophil to lymphocyte ratio

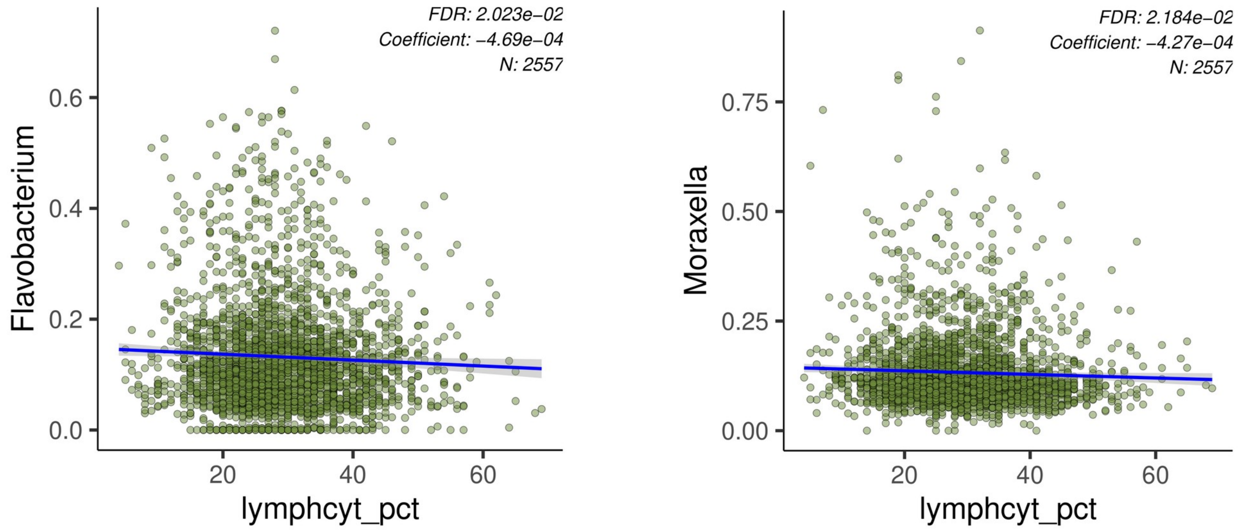

Figures E8 and E9. Plots of inferred taxa abundance and lymphocyte percentage

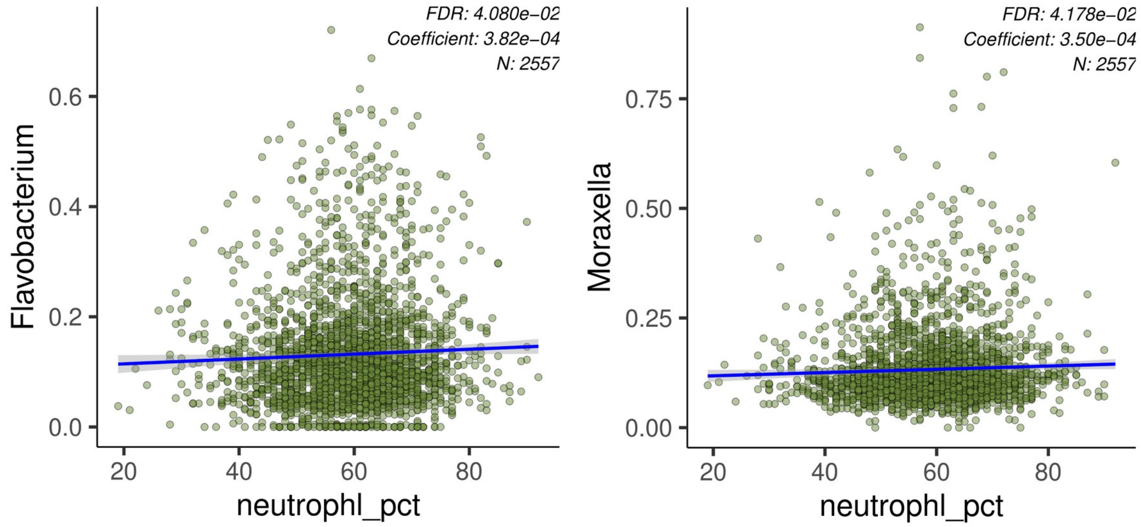

Figures E10 and E11. Plots of inferred taxa abundance and neutrophil percentage

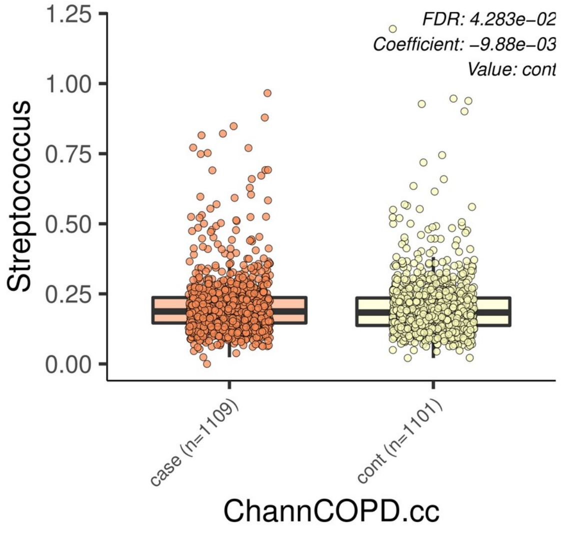

Figure E12. Plot of inferred taxa abundance and COPD case-control status

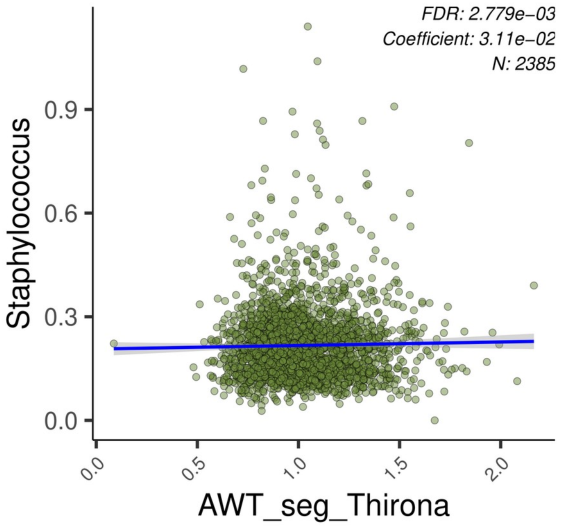

Figure E13. Plot of inferred taxa abundance and airway wall thickness

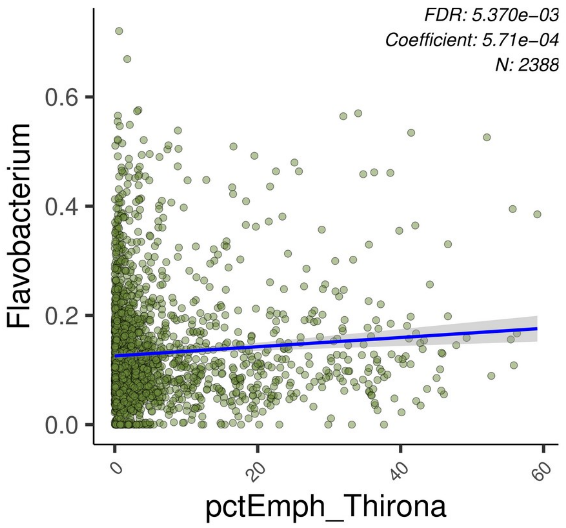

Figure E14. Plot of inferred taxa abundance and percent emphysema

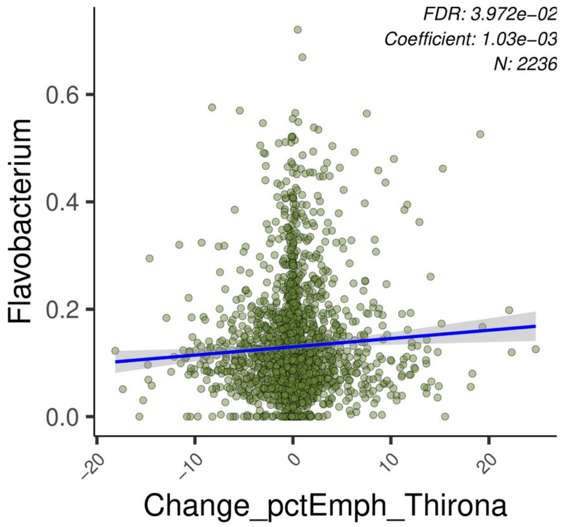

Figure E15. Plot of inferred taxa abundance and change in percent emphysema

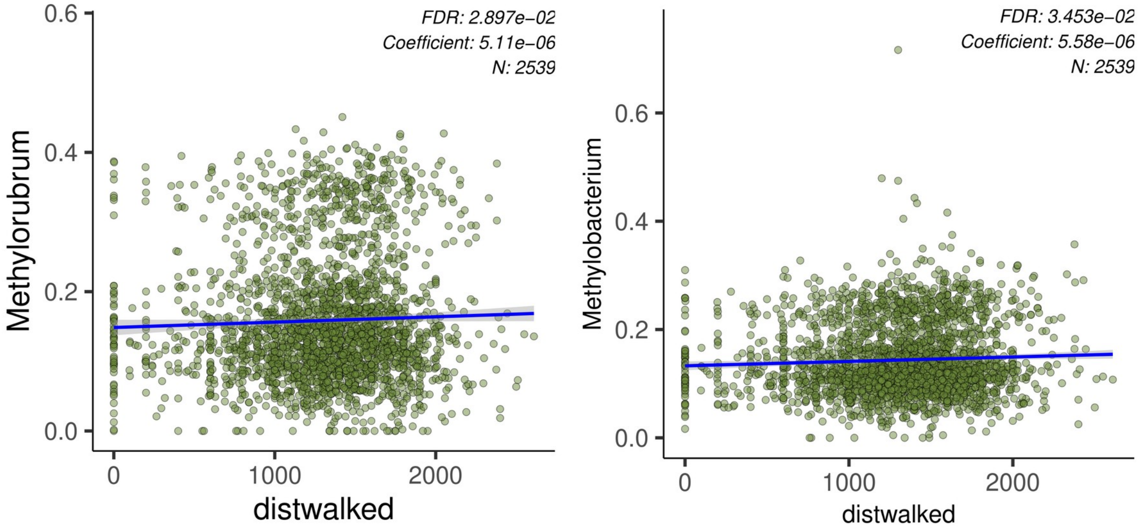

Figures E16 and E17. Plots of inferred taxa abundance and six minute walk distance

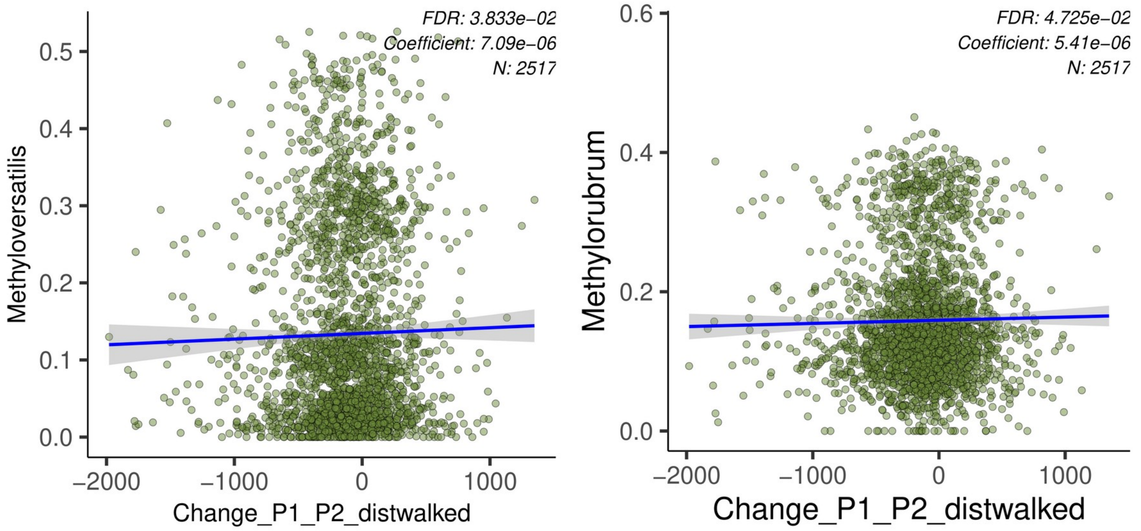

Figures E18 and E19. Plots of inferred taxa abundance and change in six minute walk distance

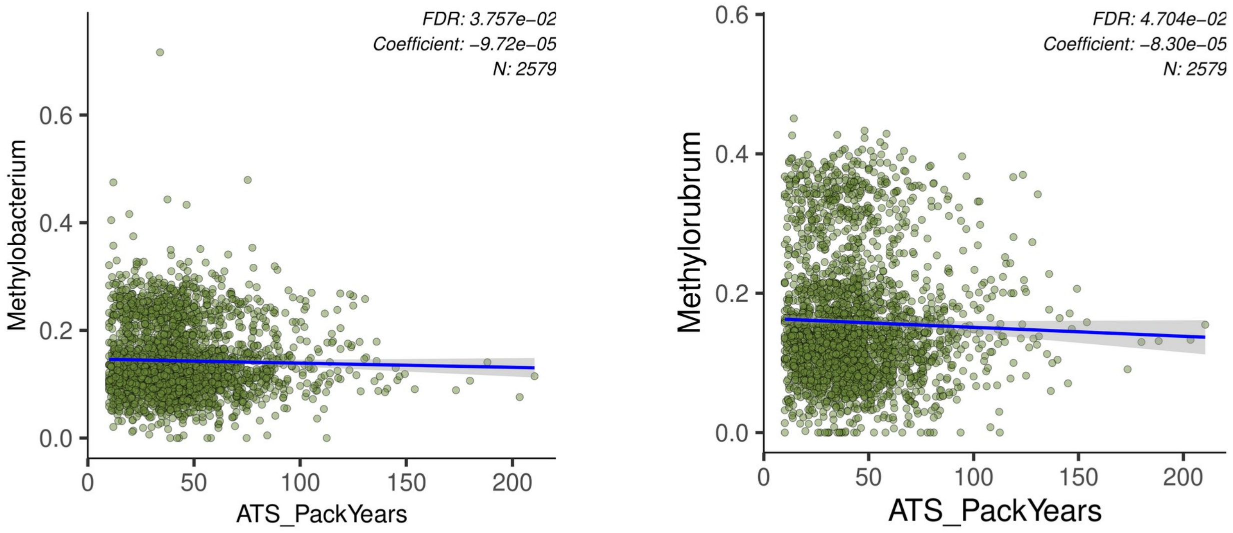

Figures E20 and E21. Plots of inferred taxa abundance and pack-years

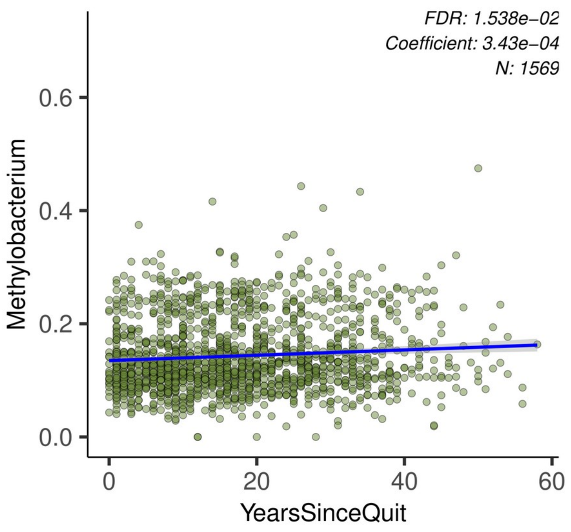

Figure E22. Plot of inferred taxa abundance and years since quit smoking

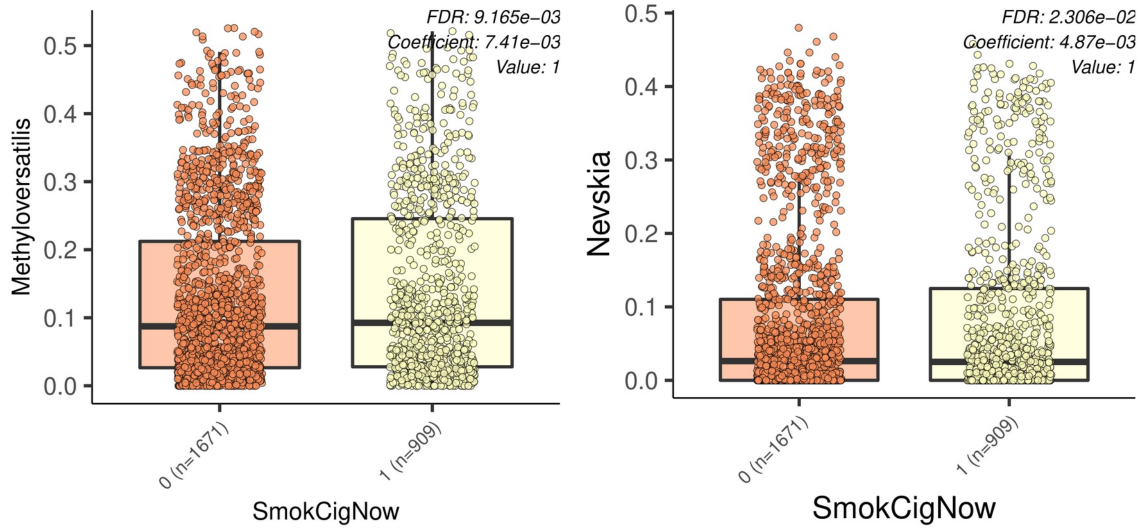

Figures E23 and E24. Plots of inferred taxa abundance and current smoking status

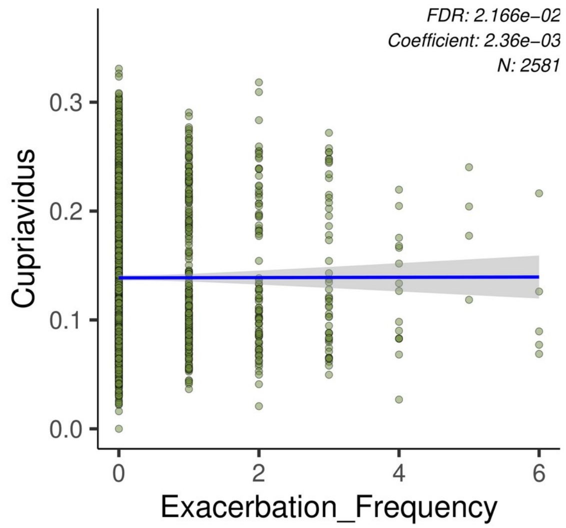

Figure E25. Plot of inferred taxa abundance and exacerbation frequency

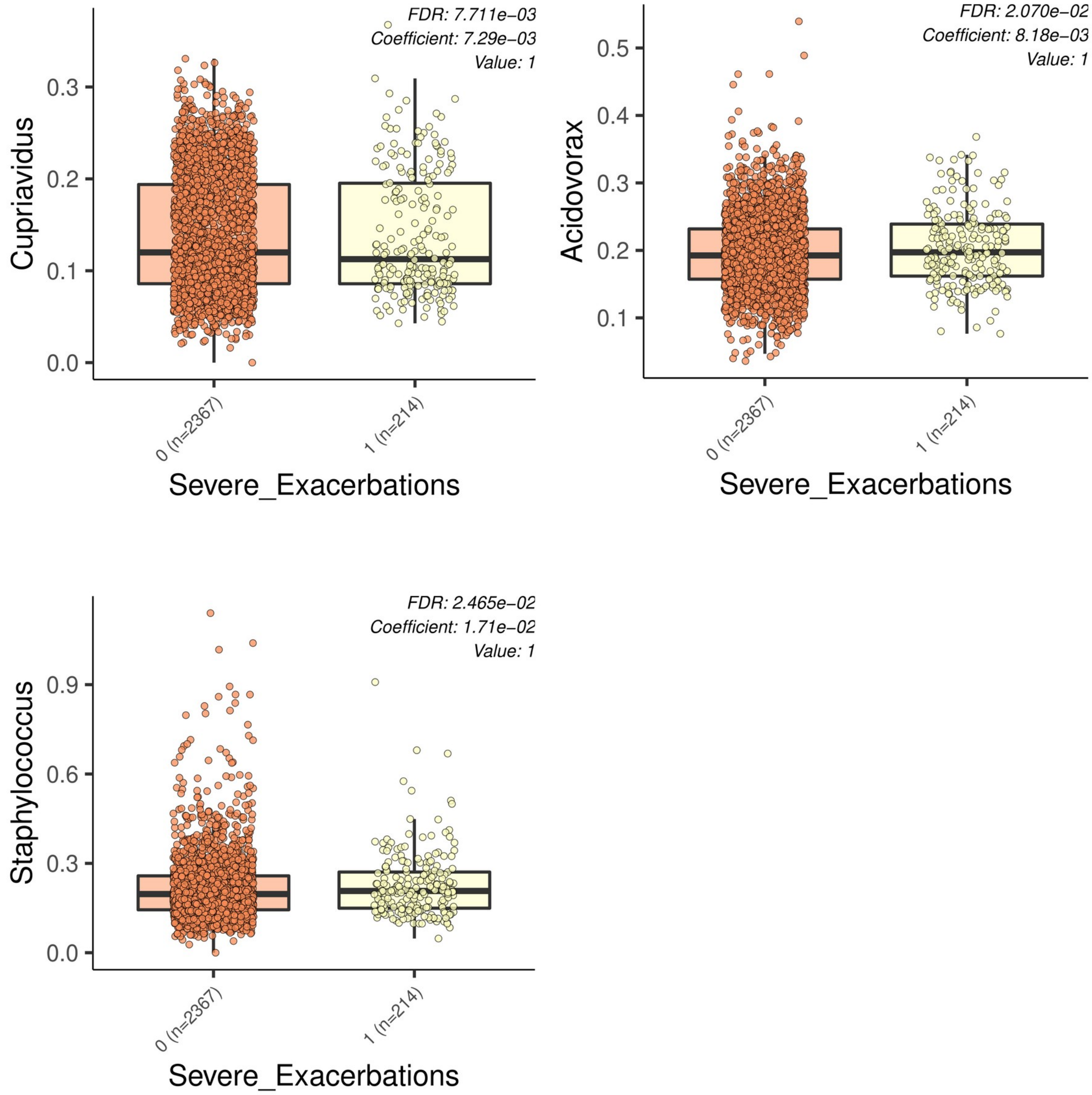

Figures E26 and E28. Plots of inferred taxa abundance and severe exacerbations

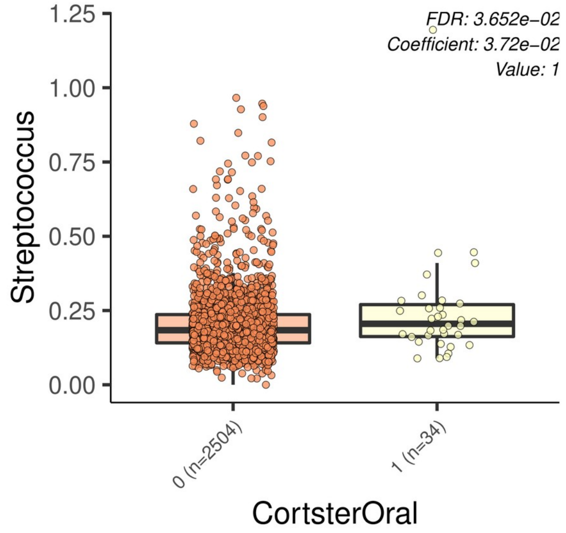

Figure E29.  Plot of inferred taxa abundance and treatment with oral corticosteroids

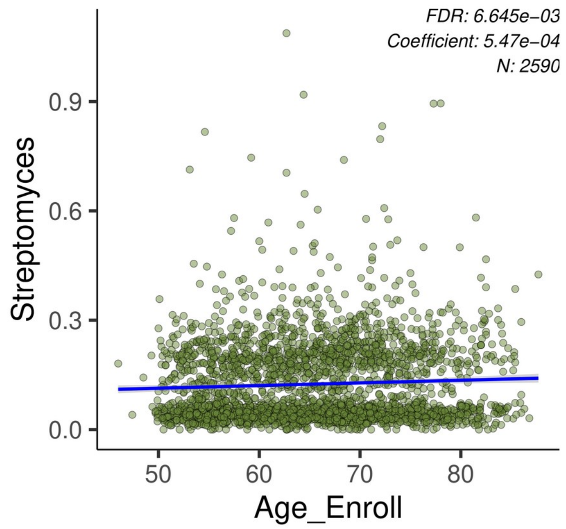

Figure E30.  Plot of inferred taxa abundance and age

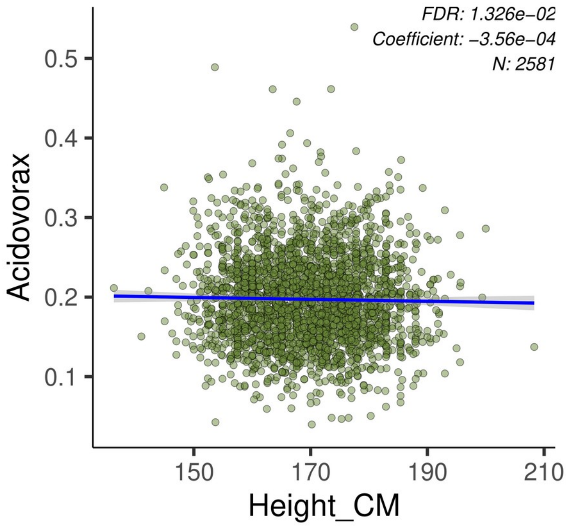

Figure E31. Plot of inferred taxa abundance and height

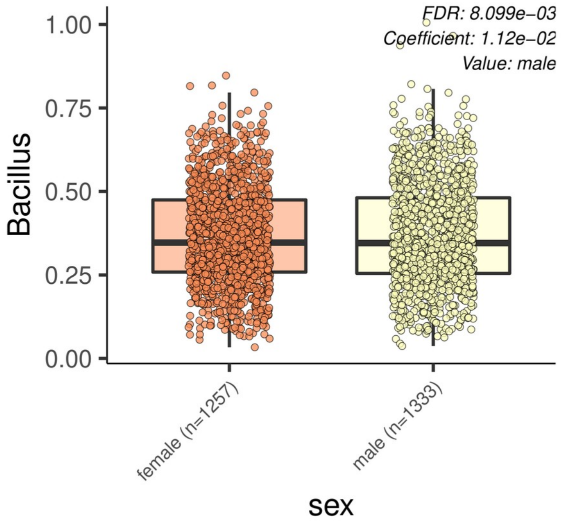

Figure E32. Plot of inferred taxa abundance and sex

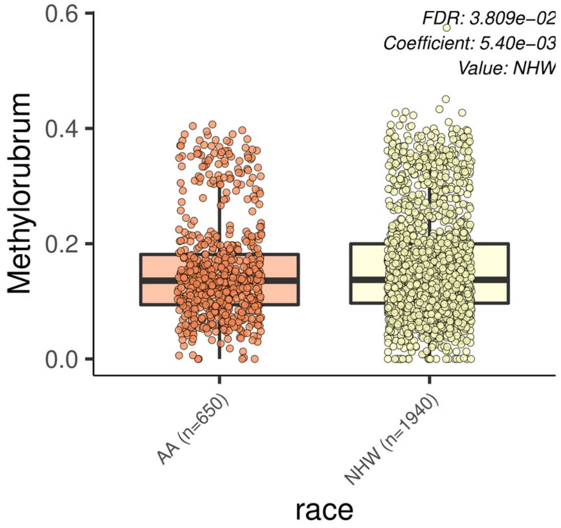

Figure E33. Plot of inferred taxa abundance and race

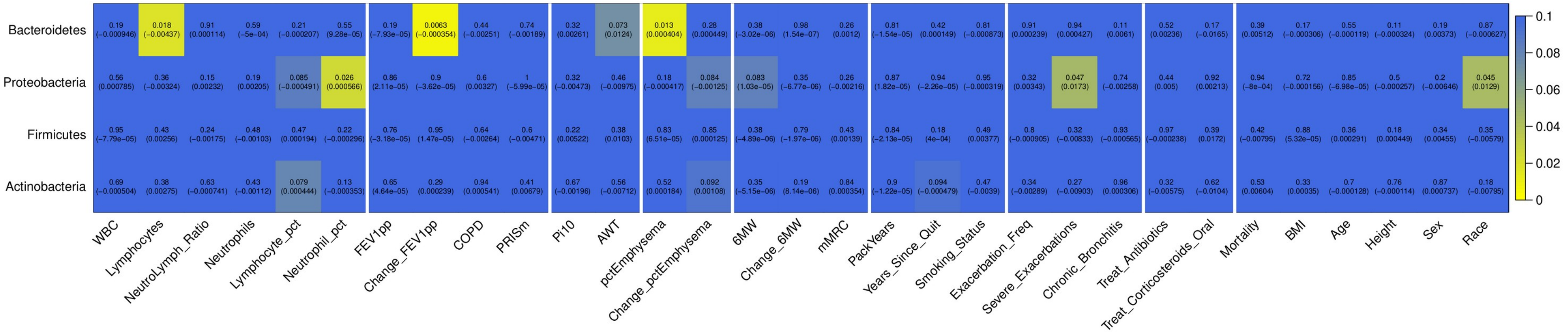

Figure E34. Heatmap of the associations between phyla inferred relative abundance and host-related variables. Top entry in each cell is the q-value (FDR adjusted p-value for each variable) and the bottom is the effect estimate from the MaAsLin2 model. Significant findings have q-values less than 0.05.

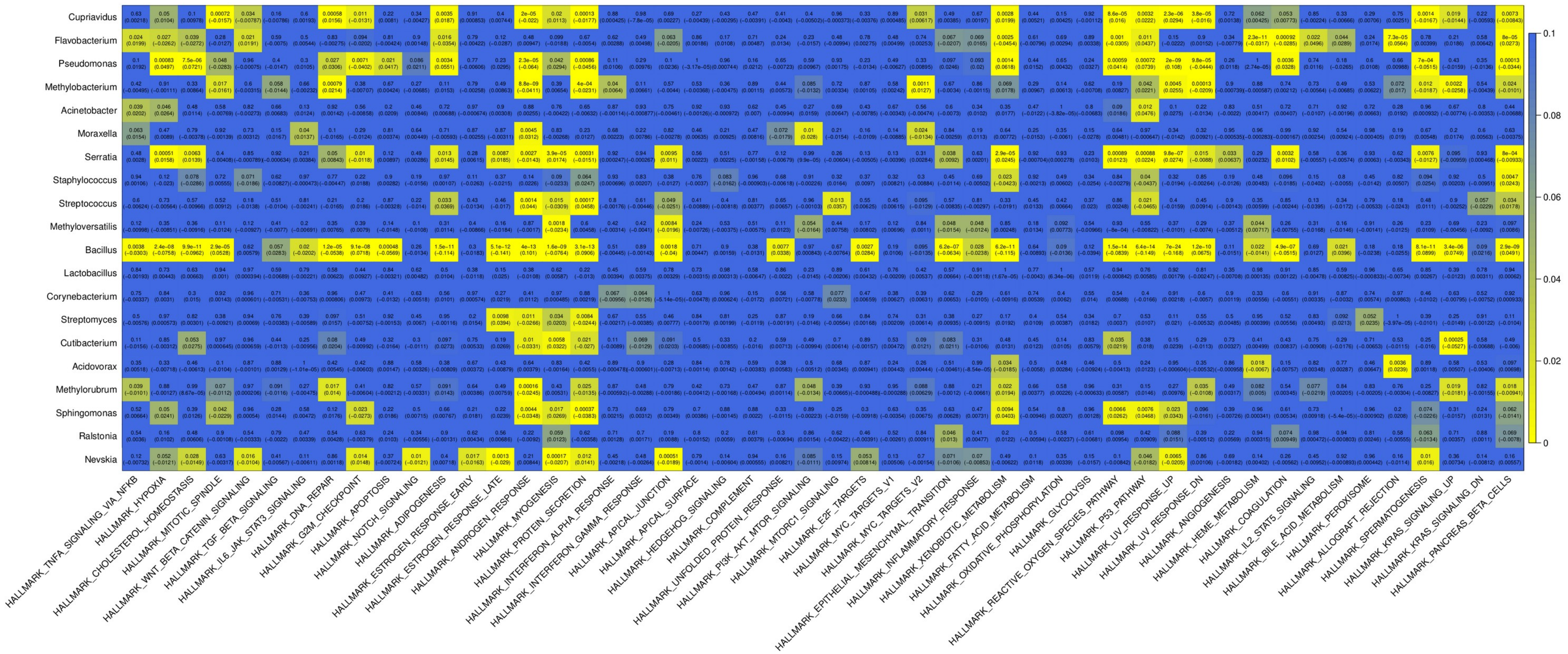

Figure E35. Heatmap of the associations between genera inferred relative abundance and Hallmark host pathways. Top entry in each cell is the q-value (FDR adjusted p-value for each variable) and the bottom is the effect estimate from the MaAsLin2 model. Significant findings have q-values less than 0.05.

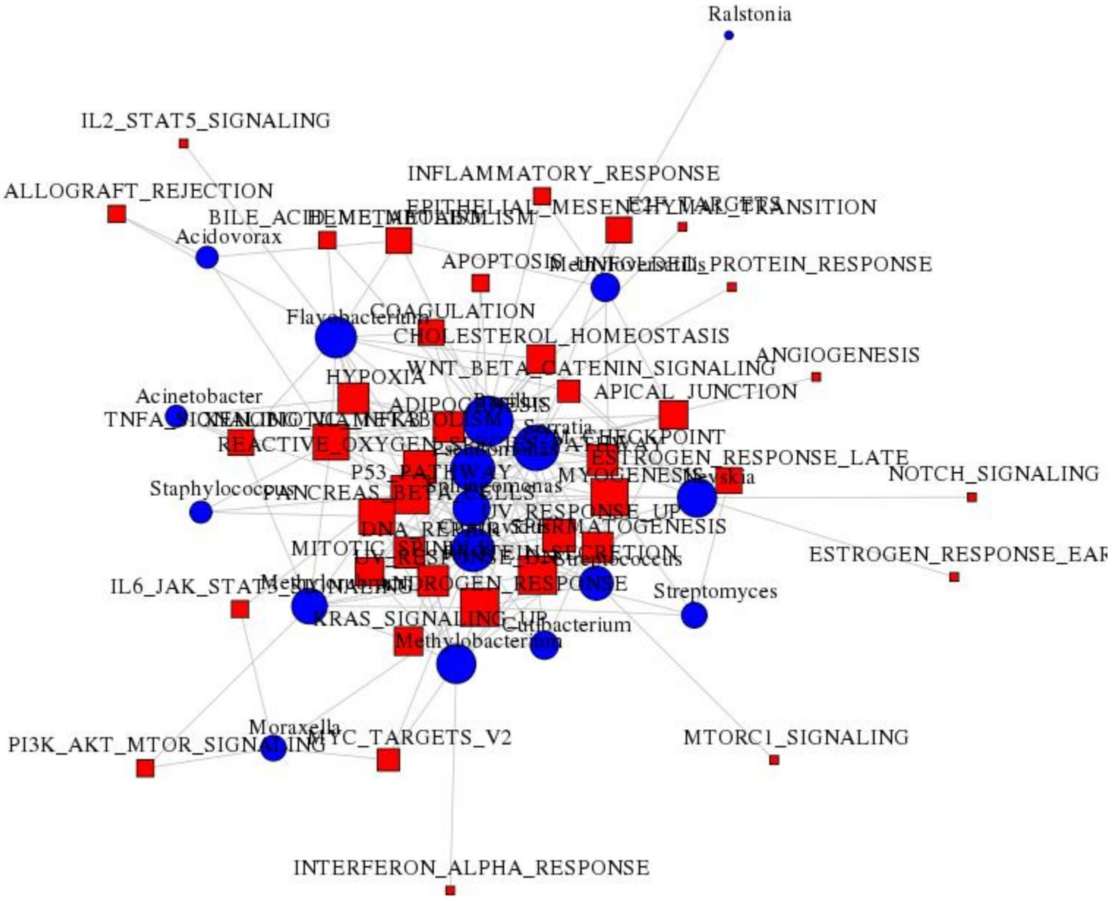

Figure E36. Bipartite network from the host-microbiome interaction analysis. Edges represent a significant (FDR < 5%) association between inferred genus abundance and the expression of the Hallmark pathway in the human host. The red squares represent Hallmark pathways from MSigDB and the blue circles represent genera.

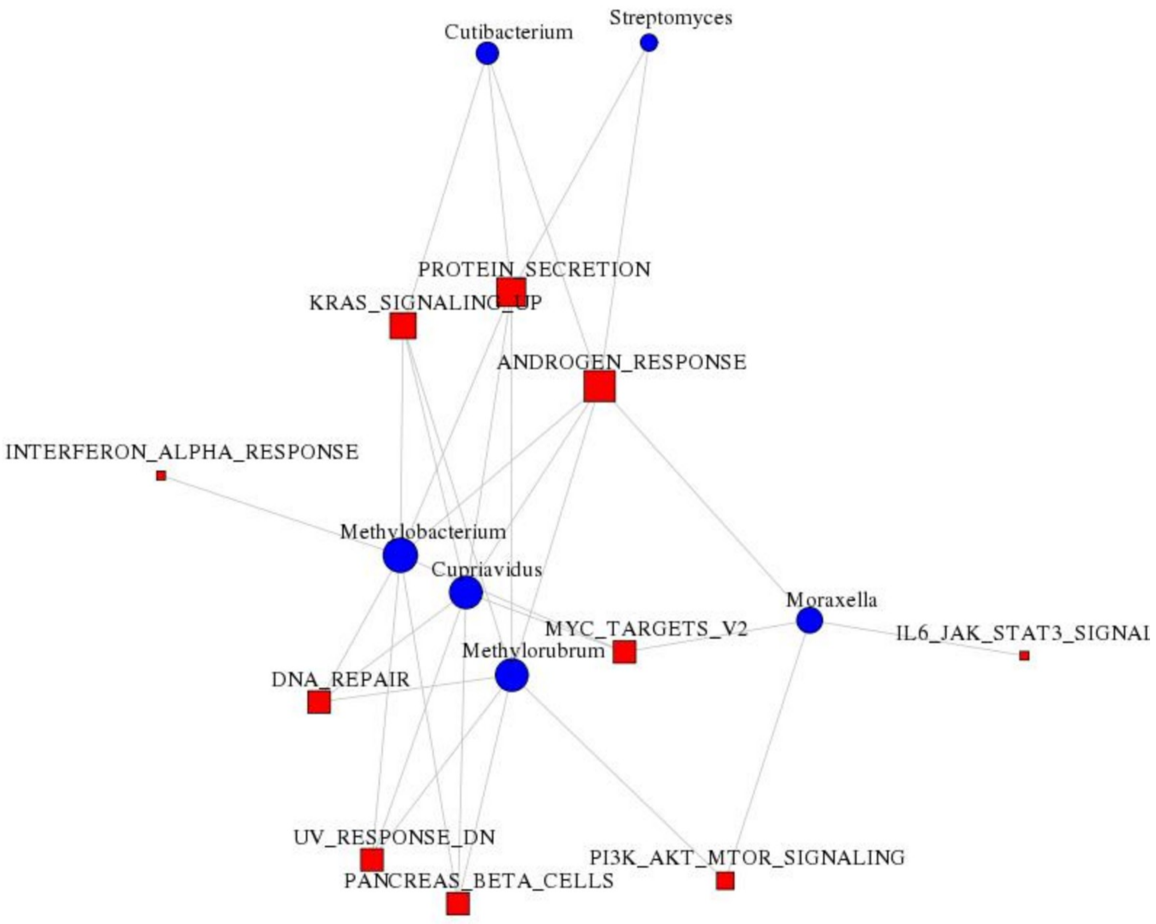

Figure E37. Community from the bipartite network from the host-microbiome interaction analysis. Edges represent a significant (FDR < 5%) association between inferred genus abundance and the expression of the Hallmark pathway in the human host. The red squares represent Hallmark pathways from MSigDB and the blue circles represent genera.

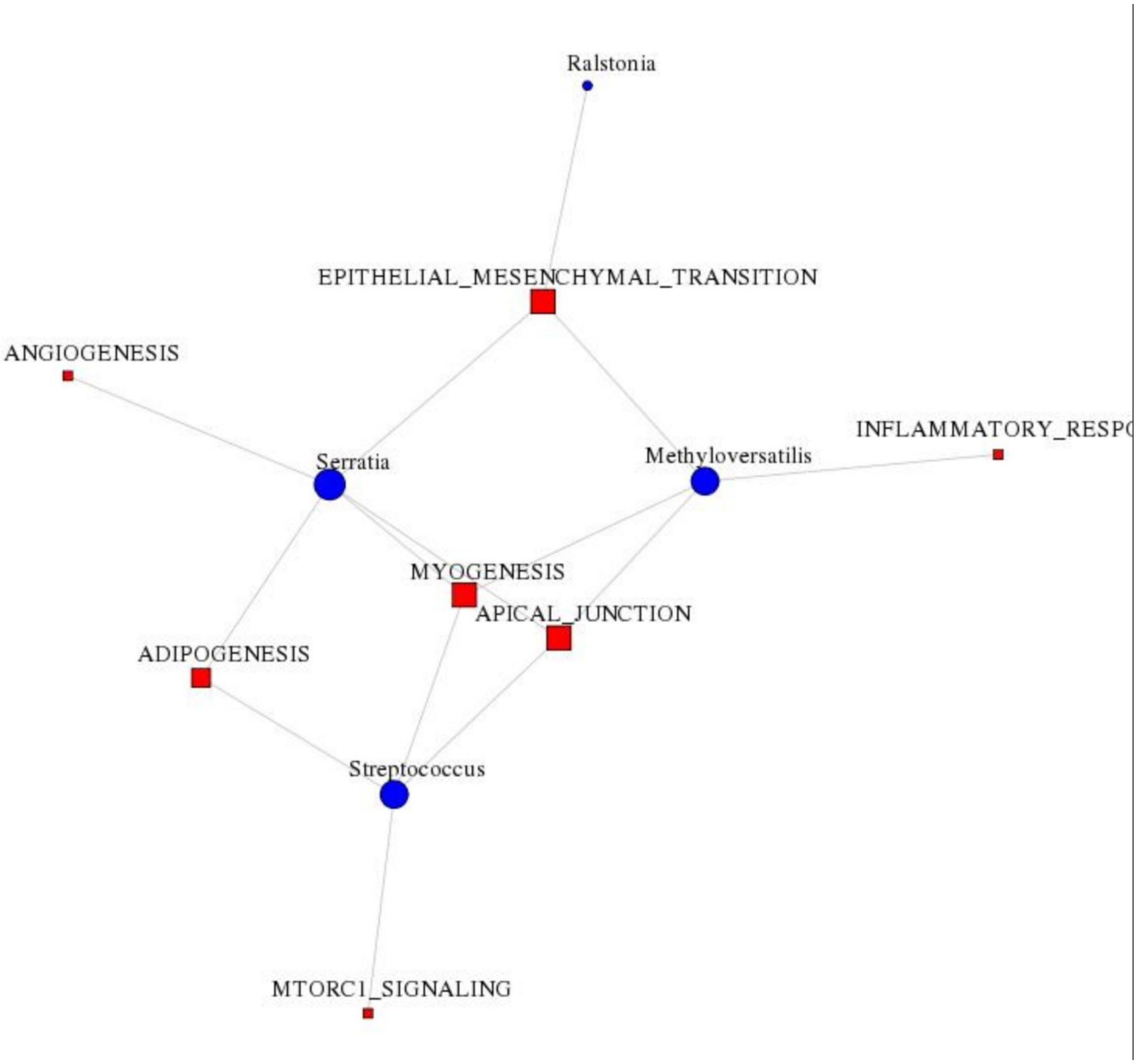

Figure E38. Community from the bipartite network from the host-microbiome interaction analysis. Edges represent a significant (FDR < 5%) association between inferred genus abundance and the expression of the Hallmark pathway in the human host. The red squares represent Hallmark pathways from MSigDB and the blue circles represent genera.

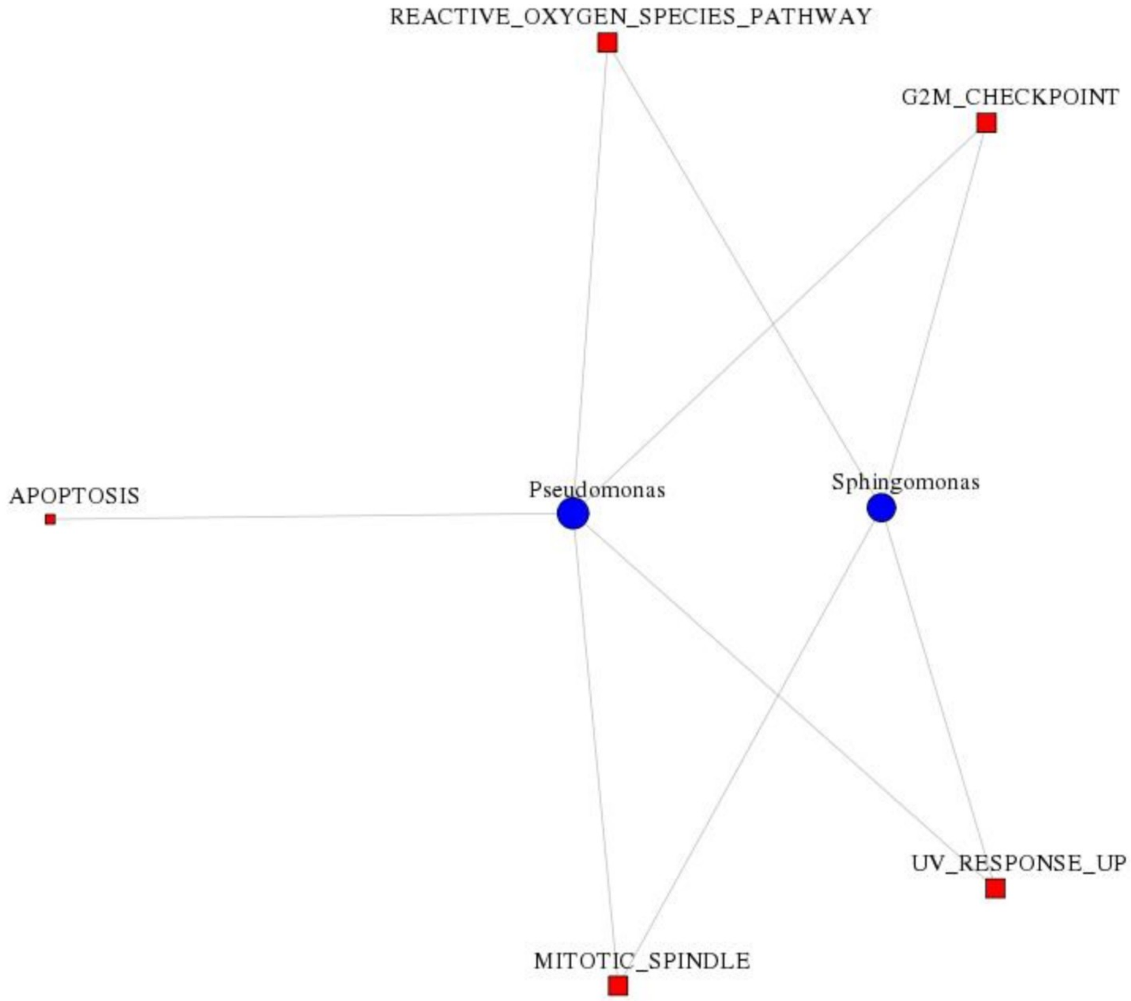

Figure E39. Community from the bipartite network from the host-microbiome interaction analysis. Edges represent a significant (FDR < 5%) association between inferred genus abundance and the expression of the Hallmark pathway in the human host. The red squares represent Hallmark pathways from MSigDB and the blue circles represent genera.

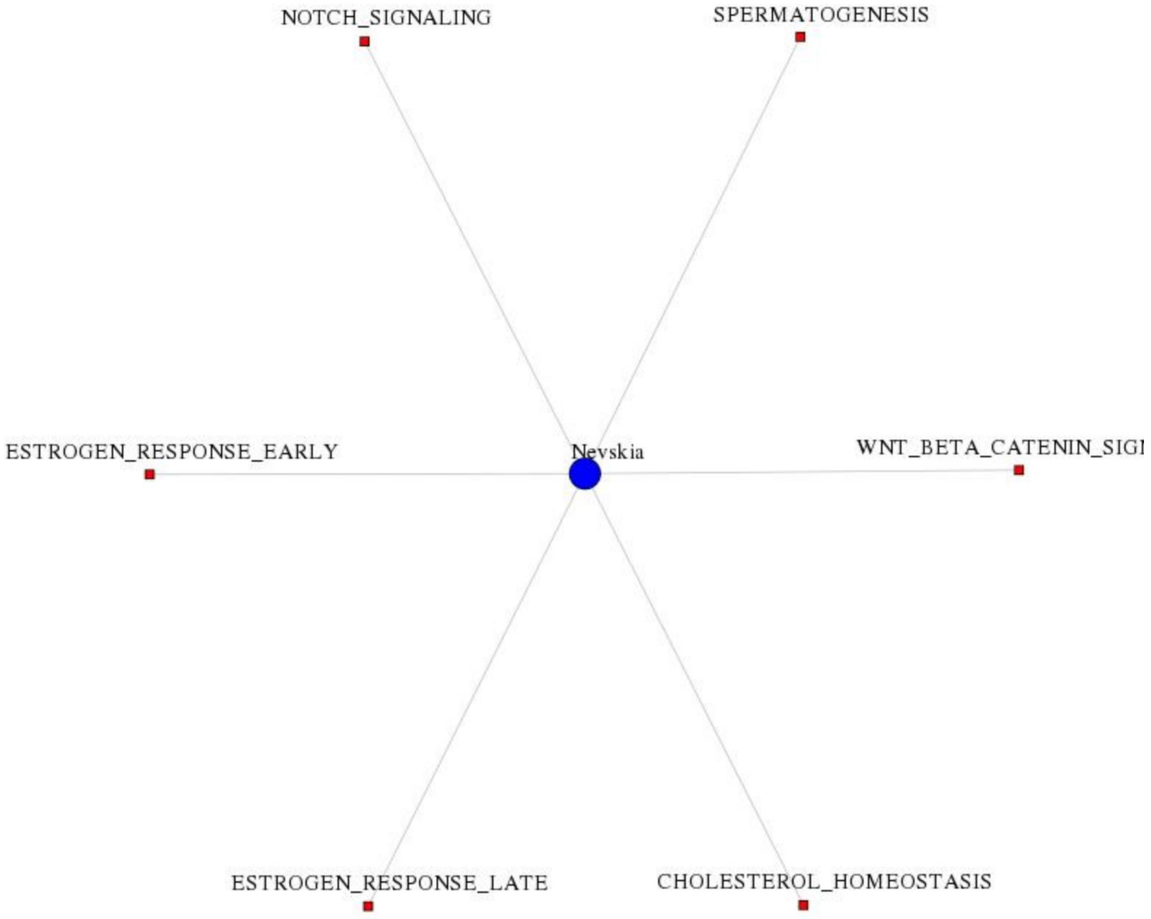

Figure E40. Community from the bipartite network from the host-microbiome interaction analysis. Edges represent a significant (FDR < 5%) association between inferred genus abundance and the expression of the Hallmark pathway in the human host. The red squares represent Hallmark pathways from MSigDB and the blue circles represent genera.

### Supplemental Tables

Table E1. Models for microbial taxon associations with outcomes of interest (inferred taxonomic abundance is the outcome in each model)

| **Predictor variable of interest** | **Model** |
|---|---|
| White blood cell count | ~ WBC + Age + Sex + Race + Smoking_Status + PackYears + batch |
| Lymphocyte count | ~ Lymphocytes + Age + Sex + Race + Smoking_Status + PackYears + batch |
| Neutrophil to lymphocytes ratio | ~ NeutroLymph_Ratio + Age + Sex + Race + Smoking_Status + PackYears + batch |
| Neutrophil count | ~ Neutrophils + Age + Sex + Race + Smoking_Status + PackYears + batch |
| Lymphocyte percentage | ~ Lymphocyte_pct + Age + Sex + Race + Smoking_Status + PackYears + batch |
| Neutrophil percentage | ~ Neutrophil_pct + Age + Sex + Race + Smoking_Status + PackYears + batch |
| FEV1 % predicted | ~ FEV1pp + Age + Sex + Race + Smoking_Status + PackYears + batch |
| Change in FEV1 % predicted [#] | ~ Change_FEV1pp + Age + Sex + Race + Smoking_Status + PackYears + batch |
| COPD: case-control [*] | ~ COPD + Age + Sex + Race + Smoking_Status + PackYears + batch |
| COPD: prism-control [@] | ~ PRISm + Age + Sex + Race + Smoking_Status + PackYears + batch |
| Pi10 | ~ Pi10 + Age + Sex + Race + Smoking_Status + PackYears + batch |
| Airway Wall Thickness | ~ AWT + Age + Sex + Race + Smoking_Status + PackYears + batch |
| Percent Emphysema | ~ pctEmphysema + Age + Sex + Race + Smoking_Status + BMI + PackYears + batch |
| Change in Percent Emphysema [#] | ~ Change_pctEmphysema + Age + Sex + Race + Smoking_Status + BMI + PackYears + batch |
| 6-minute walk distance (ft) | ~ 6MW + Age + Sex + Race + Smoking_Status + PackYears + batch |
| Change in 6-minute walk distance (ft) [#] | ~ Change_6MW + Age + Sex + Race + Smoking_Status + PackYears + batch |
| MMRC dyspnea score | ~ mMRC + Age + Sex + Race + Smoking_Status + PackYears + batch |
| Pack years of smoking | ~ PackYears + Age + Sex + Race + Smoking_Status + batch |
| Years since quit smoking | ~ Years_Since_Quit + Age + Sex + Race + PackYears + batch |
| Smoking status | ~ Smoking_Status + Age + Sex + Race + PackYears + batch |
| Exacerbation Frequency | ~ Exacerbation_Frequency + Age + Sex + Race + Smoking_Status + PackYears + batch |
| Severe Exacerbations (yes / no) | ~ Severe_Exacerbations + Age + Sex + Race + Smoking_Status + PackYears + batch |
| Chronic Bronchitis (yes / no) | ~ Chronic_Bronchitis + Age + Sex + Race + Smoking_Status + PackYears + batch |
| Treated with antibiotics (yes / no) | ~ Treat_Antibiotics + Age + Sex + Race + Smoking_Status + PackYears + batch |
| Treated with oral corticosteroids (yes / no) | ~ Treat_Corticosteroids_Oral + Age + Sex + Race + Smoking_Status + PackYears + batch |
| Status (alive / diseased) | ~ Mortality + Age + Sex + Race + Smoking_Status + PackYears + batch |
| Body mass index (kg/$m^2$) | ~ BMI + Age + Sex + Race + Smoking_Status + PackYears + batch |
| Age (years) | ~ Age + Sex + Race + Smoking_Status + PackYears + batch |
| Height (cm) | ~ Height_CM + Age + Sex + Race + Smoking_Status + PackYears + batch |
| Sex | ~ Sex + Age + Race + Smoking_Status + PackYears + batch |
| Race | ~ Race + Age + Sex + Smoking_Status + PackYears + batch |

| |
|---|
| Smoking_Status (ordinal variable): 0 former smoker, 1 current smoker |
| Abbreviations: FEV1=forced expiratory volume in 1 sec; pctEmph=% emphysema; Pi10=SRWA-Pi10=square root wall area of a hypothetical airway with 10mm internal perimeter; AWT=airway wall thickness; batch=processing batch variable;<br><br>* PRISm subjects (GOLD = -1 excluded)<br>@ COPD cases (GOLD = 1,2,3,4 excluded)<br># Change variables reflect COPDGene Phase 1 visit to Phase 2 visit |

Tables E2 through E32. Results from tests of association between the inferred relative abundance for each taxon at the genus level and host phenotype, exposure, treatment and trait variables using linear models with MaaAsLin2.

See file: Tables_E2_to_E32.xls
